# Romaciclib, a CDK8/CDK19 inhibitor, can overcome venetoclax resistance through a combinatorial strategy

**DOI:** 10.64898/2025.12.16.693978

**Authors:** Urszula Pakulska, Marta Obacz, Jerzy Woźnicki, Katarzyna Wiklik, Samarpana Chakraborty, Marta Micek, Vakul Mohanty, Diego Coelho, Elżbieta Adamczyk, Aniela Gołas, Adrianna Moszyńska, Hui Zhang, Magdalena Cybulska-Lubak, Ewelina Kaniuga, Zuzanna Sandowska-Markiewicz, Marina Konopleva, Michał Mikula, Przemysław Juszczyński, Aditi Shastri, Natalia Baran, Tomasz Rzymski, Milena Mazan

## Abstract

The combination of venetoclax (VEN) and hypomethylating agents (HMA) is the standard of care in acute myeloid leukemia (AML) for elderly patients unfit for intensive chemotherapy. Despite its clinical success, most patients eventually relapse, creating an urgent need for effective therapeutic alternatives. In this study, we aimed to evaluate the potential of romaciclib, a first-in-class CDK8/CDK19 inhibitor, in combination with VEN to overcome stroma-mediated and primary/acquired VEN-resistance. We assessed the efficacy of RVU120+VEN combination in both sensitive and resistant AML cell lines and primary patient-derived models. Our finding demonstrated that romaciclib synergizes with VEN in AML cell lines and in 8 out of 11 patient-derived cell samples. The proteomic and functional studies demonstrated that combination induced apoptosis through caspase-dependent cleavage of MCL-1. *In vivo* studies confirmed the efficacy of RVU120+VEN, showing eradication of leukemic cells and bone marrow recovery. Importantly, the combination effectively overcame both stroma-mediated and transcriptionally dependent VEN-resistance. Mechanistic studies, focusing on transcriptomic analyses, identified key resistance-associated pathways, including IL6/JAK/STAT3, TGF-β, PI3K/AKT/MTOR, and inflammatory signaling, being suppressed by combination treatment. Furthermore, an in vivo study using a VEN-resistant patient-derived xenograft (PDX) model confirmed the efficacy of the combination, demonstrating a significant reduction in leukemia burden and a decreased proportion of leukemia initiating cells (LIC) following treatment. These findings prove the highly synergistic mechanism of action of RVU120+VEN combination and the potential to overcome primary/acquired VEN resistance in relapse/refractory AML disease. Altogether, the presented results support ongoing clinical studies evaluating romaciclib and VEN in VEN/HMA-refractory patients (NCT06191263) and provide a basis for future exploration as a frontline therapy in VEN-naïve patients.

## Introduction

Acute Myeloid Leukemia (AML) is characterized by the accumulation of immature myeloid blasts in the bone marrow (BM) and peripheral blood (PB), impairing hematopoiesis and causing cytopenia.^1^ It mainly affects elderly patients,^2^ many of whom are unfit for intensive chemotherapy. For this group, venetoclax (VEN) combined with azacitidine (AZA) has shown strong efficacy, leading to US Food and Drug Administration approval as a standard treatment.^3,4^

VEN is a selective inhibitor of the anti-apoptotic B-cell lymphoma 2 (BCL-2) protein that promotes mitochondrial apoptosis in AML cells.^5^ However, upregulation of alternative anti-apoptotic proteins, such as myeloid leukemia 1 (MCL-1), can diminish its efficacy. Dual targeting of both BCL-2 and MCL-1 enhances apoptosis in both VEN-sensitive and -resistant AML, though clinical safety remains a concern.^6^

Effective treatment options for AML patients who are refractory or relapse (R/R) after VEN plus hypomethylating agents are critically needed.^7,8^ These patients face poor outcomes due to the complex, heterogeneous nature of VEN resistance, limited salvage therapies, and low clinical trial access.^9^

Beyond BCL-2 family proteins resistance is also linked to AML differentiation, metabolic rewiring, and mitochondrial adaptations^10–14^. Recent studies underscore the intra- and inter-patient heterogeneity of VEN resistance; for example, Mohanty et al identified distinct transcriptional clusters associated with diverse pathways of resistance to VEN^14^.

The BM microenvironment plays a pivotal role in the development of drug-resistance. Stromal cells protect leukemic cells from cytotoxic drugs through direct cell-to-cell interactions, secretion of soluble factors, and modulation of cell-intrinsic survival pathways.^15–17^ Stromal cells also contribute to kinase inhibitor resistance, with early BM-driven resistance evolving into mutation-dependent late-stage resistance.^18^

In the landscape of AML treatments, Cyclin-Dependent Kinase 8 (CDK8) has emerged as a promising therapeutic target. Romaciclib (RVU120, SEL120-34A), a potent first-in-class CDK8/CDK19 inhibitor, is currently in phase II clinical trials for various hematologic malignancies. Preclinical studies have demonstrated significant antileukemic activity of romaciclib and other CDK8 inhibitors across multiple AML models *in vitro* and *in vivo*.^19,20^ In this study, we demonstrate that the combination of romaciclib and VEN exhibits strong synergy in AML cell lines and patient-derived cells, overcoming multiple mechanisms of VEN resistance. Specifically, RVU120+VEN downregulates the anti-apoptotic protein MCL-1, counteracts stromal-mediated resistance, and shows superior efficacy in AML cells from two of the four recently characterized transcriptionally distinct VEN-resistant clusters.^14^ Mechanistically, RVU120+VEN treatment suppressed key resistance-associated inflammatory signaling in AML cells. Notably, this combination is being further investigated in the RIVER81 study (ClinicalTrials.gov ID NCT06191263), underscoring its potential as a novel therapeutic strategy for overcoming VEN resistance in AML.

## Materials and Methods

### Cell lines and inhibitors

AML cell lines and HS-5 mesenchymal stromal cells were cultured in their recommended media (**Supplemental Table S1**) at 37⁰C in a humidified atmosphere containing 5% CO_2_. All cell lines were authenticated using short tandem repeat profiling, and the absence of mycoplasma was confirmed by the MycoAlert assay (Lonza LT07-518). Romaciclib was provided by Ryvu Therapeutics. VEN, Z-VAD-FMK, MG-132, and Staurosporine were obtained from MedChemExpress, ChemScene, or AA Blocks.

### Patient-derived primary cells and patient-derived xenograft culture

A patient-derived xenograft (PDX) AMLX12 model was cultured for up to 72 hours in its recommended medium (**Supplemental Table S1**). PB from AML was obtained from patients with informed consent, according to the requirements of MD Anderson Cancer Center Institutional Review Boards (LAB PA13-1025) and the Declaration of Helsinki. Primary patient derived AML cells (PDCs) isolation was performed as previously described^21,22^ and followed by incubation with selected compounds for 96 hours. The characteristics of primary samples are summarized in **Supplemental Table S2**.

### Viability assays, IC50 and synergy calculation

Cell lines were seeded at density range 7000-40,000 cells/well in 96-well plates, and PDX cells at 15,000 cells/well in 384-well plates. Cells were exposed to romaciclib and/or VEN for 72 or 96 hours. For long-term viability studies, on days 3, 7, 10, and 14, an equal volume of cell suspension was transferred into fresh drug-containing media to restore the initial seeding density for Dimethyl sulfoxide controls. Cell viability was measured on the indicated days using CellTiter-Glo assy (CTG) at a 1:3 ratio (Promega G9242) or AlamarBlue at a 1:20 ratio (Thermo Scientific DAL1100) using the EnSpire microplate reader (PerkinElmer). IC50s were calculated using a nonlinear regression model in GraphPad Prism (v.10.1.2). Drug synergy was assessed using the Loewe additivity model in SynergyFinder Plus software.

### Apoptosis assays

Cell lines were treated with romaciclib and/or VEN for 24 hours or 96 hours and analyzed by flow cytometry using an Annexin V antibody (1:25, BD 556421 or 556419) and 7AAD (1:25, BD 559925) on a BD LSR Fortessa cytometer. Flow cytometry analysis was performed using FlowJo software (v.10.8.1). PDCs from AML patients (n = 11) were seeded at 300,000 cells/well in 24-well plates and treated as detailed in the figure legend. The evaluation of specific apoptosis was calculated as previously described.^21,22^

### Stromal cell-conditioned medium

To generate conditioned media, 3×10^6^ of HS-5 stromal cells were plated in 10 ml of culture medium for 72 hours, then media were harvested and cleared by centrifugation. MOLM-14 and MV4-11 cell lines were seeded at 10,000 cells/well into 96-well plates in a 1:1 mixture of HS-5-conditioned media and their standard culture media. Cells were exposed to romaciclib and VEN for 72 hours. Viability, IC50s, and drug synergy were calculated as described above.

### Western blotting

Cells were lysed in a RIPA buffer (Thermo Scientific 89900) supplemented with phosphatase and protease inhibitors (Thermo Scientific 78444). The lysates were clarified by centrifugation and quantified using Bradford assay (Sigma B6916). Equal amounts of whole-cell lysates were loaded on 4-15% tris-glycine gels (Bio-Rad 5678085), then transferred to PVDF membranes. Immunoblotting was performed using antibodies listed in **Supplemental Table S3**. Proteins were visualized using the ChemiDoc MP Imaging System (Bio-Rad). Densitometry was performed using the ImageLab software (v.5.1).

### RNA-Seq processing and data analysis

MV4-11, MOLM-13, KG-1, and PDX-AMLX12 cells were treated with romaciclib and/or VEN in biological triplicates/condition as described in the figure legends. Total RNA was isolated using NucleoSpin RNA Mini kit (Macherey-Nagel 740955). RNA-Seq data were processed and analyzed as detailed in the **Supplemental Methods**.

### AML orthotopic xenograft study

For studies of efficacy of RVU120+VEN *in vivo*, cell line–derived xenograft (CDX) models were applied. The animal welfare complied with the UK Animal Scientific Procedures Act 1986 in line with Directive 2010/63/EU. We injected 84 female NOD SCID gamma (NSG) mice, 6-8 weeks old, intravenously with 5 mln MV4-11 (ATCC) Luc-tagged cells. Based on bioluminescence imaging (BLI) on day 8, mice were randomized into six experimental groups (n = 10): vehicle control, romaciclib at 20 or 40 mg/kg, VEN at 100 mg/kg, or romaciclib (both doses) plus VEN combination groups. From day 9, drugs were administered once daily by oral gavage. Body weight (BW) was measured daily and the leukemic burden was monitored by BLI once a week. After day 21, mice were sacrificed over three consecutive days. BM was isolated and analyzed by flow cytometry for hCD45 and mCD45, as detailed in the **Supplemental Methods**.

### AML subcutaneous xenograft study

The study was performed according to the protocol approved by the Institutional Animal Care and Use Committee of Shanghai ChemPartner following the guidance of the American Association for Accreditation of Laboratory Animal Care. We injected 90 female BALB/c nude mice, 6-9 weeks old, subcutaneously with MV4-11 (ATCC) cells. BW and tumor volume were measured three times per week. On day 14, mice were randomly distributed into six treatment groups (n = 10): vehicle control, romaciclib at 45 mg/kg, VEN at 100 mg/kg, or combinations of romaciclib at 15, 30, and 45 mg/kg with VEN (100 mg/kg). Drugs were administered once daily by oral gavage. The study was terminated during day 21-22 of dosing. Details in the **Supplemental Methods**.

### AML patient derived xenograft study

The study was performed under protocols approved by the institutional animal facility at Albert Einstein College of Medicine. NOD-scid Il2rg-KO NSG mice were purchased from Jackson Laboratory and maintained under pathogen-free conditions and monitored daily. Human VEN-resistant AML patient-derived BM cells (PDX-S1) were thawed, washed in PBS supplemented with 2% FBS, and resuspended in sterile PBS. The mice were sublethally irradiated (2.5Gy) 24 hours before injection and injected with 1 × 10^5 cells per mouse intravenously via the tail vein. The engraftment was confirmed 5 weeks post-transplantation (>0.3% hCD45+ cells in BM aspirate) and animals were randomized into four treatment groups (n=10 per group): vehicle control, romaciclib at 20 mg/kg, VEN at 100 mg/kg, or combinations of romaciclib 20 mg/kg with VEN 100 mg/kg. Dosing scheme and formulation details are described in the **Supplemental Methods.** Following four weeks of treatment, mice were euthanized. BM and spleen cells were harvested, frozen and stored in -80°C.

### Flow cytometry analysis of PDX samples

AMLX-12 cells were incubated with Human Fc Block (BD 564220) and anti-mouse CD16/CD32 antibody (Invitrogen 14016186) to reduce nonspecific antibody staining caused by IgG receptors. Following blocking, samples were stained with an anti-human antibody panel containing: CD45 BUV395 (1:40, BD 563792), CD34 PE (1:20, BioLegend 343506), CD38 PE/Dazzle594 (1:20, BioLegend 356630), CD45RA APC (1:40, BioLegend 304112), CD33 PerCP/Cy5.5 (1:20, BioLegend 366616), CD11b PE/Cy7 (1:20, BioLegend 301321), CD14 FITC (1:20, BioLegend 367116), and CD123 BV711 (1:40, BioLegend 306030). Additionally, samples were stained with LIVE/DEAD Fixable Near-IR (Invitrogen L34976) to assess cell viability. After singlet exclusion, viable cells were gated, followed by selection of hCD45 positive cells, and subsequent analysis of surface marker expression. Flow cytometry analysis was performed on a BD LSR Fortessa cytometer and using FlowJo software (v.10.8.1).

PDX-S1 BM cells were thawed and washed with PBS supplemented with 2% FBS (FACS Buffer). Cells were then stained with Zombie-NIR for live dead detection, followed by staining using antibody cocktail for 30 min with the following antibodies: mCD45 FITC (1:100 eBioscience 11-0453-85) , hCD45 APC (1:100, BioLegend 304037), hCD33 BV421(1:100, BioLegend 366622), hCD34 PE (1:100, BioLegend 343506), hCD3 PE-Cy7 (1:100, BioLegend 317334), hCD19 PerCP Cy5.5 (1:100, BioLegend 302230). After staining, cells were washed twice and resuspended in FACS buffer. Flow cytometry was performed on BD FACS Aurora instrument and analyzed using FlowJo Software (Version 10). Percentages of each population were calculated relative to total live BM cells.

### Statistical analysis

Values are presented as mean ± standard deviation (SD), unless otherwise indicated. Viability experiments using cell lines were performed at least twice in triplicate (apoptosis in monoplicate). Experiments using AMLX-12 cells or PDCs were performed once in triplicate, due to a limited sample size. In the in vivo experiment using PDX-S1, seven mice per group were analyzed. Data were analyzed for normal distribution, and correlation analysis between group testing were performed as described in the figure legends using GraphPad Prism software (v.10.1.2). P value < 0.05 was considered statistically significant: * p < 0.05, ** p < 0.01, and *** p < 0.001.

## Results

### Romaciclib synergistically enhances antileukemic activity of VEN in AML cells

To assess whether romaciclib enhances the antileukemic activity of VEN, we selected two VEN-sensitive AML cell lines, MV4-11 and MOLM-13, which exhibit relatively higher and lower sensitivity to romaciclib monotherapy (**Figure 1A**). Both cell lines were co-treated with a wide concentration range of romaciclib and VEN. As shown in **Figure 1B**, the RVU120+VEN combination significantly reduced cell viability compared to either monotherapy, revealing a strong synergistic interaction with mean Loewe synergy scores of 14.41 for MV4-11 and 9.8 for MOLM-13 cells.

**Figure 1.**
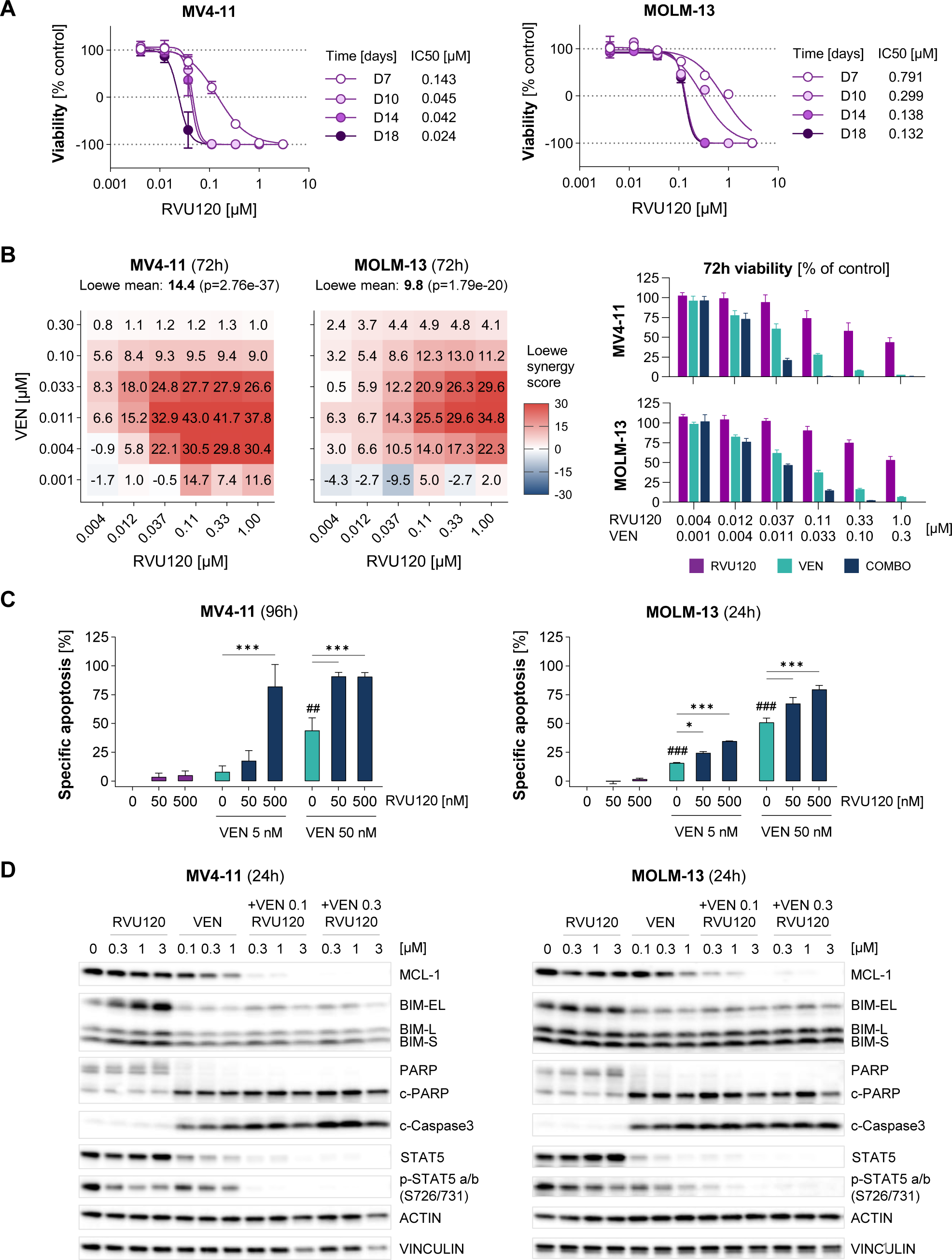
Romaciclib synergistically enhances antileukemic activity of VEN in VEN-sensitive AML cells. **(A)** Long-term viability of MV4-11 (left) and MOLM-13 (right) cells treated with increasing concentrations of romaciclib for 7, 10, 14, 18 days. Data are mean ± SD of technical triplicates from a representative experiment (n = 2). **(B)** Loewe synergy scores calculated from the effect of RVU120+VEN combination on viability in MV4-11 and MOLM-13 cells treated with the indicated romaciclib /VEN concentrations for 72 hours. Loewe scores of > 0 are regarded as likely synergistic. The bar graphs show matched cell viability at selected doses. Data are mean ± SD of technical triplicates from a representative experiment (n = 3). **(C)** Apoptosis induction in MV4-11 and MOLM-13 cells treated with romaciclib (50, 500 nM), VEN (5, 50 nM), or their combination at the indicated time points. Data are mean ± SD of two independent experiments. Two-way ANOVA with Tukey’s multiple comparisons: *p < 0.05, ***p < 0.001 (as indicated); ^##^p < 0.01, ^###^p < 0.001 (vs vehicle control). **(D)** Protein expression of MCL-1, BIM isoforms, PARP and cleaved PARP (cPARP), cleaved caspase-3 (c-caspase-3), STAT5 total and p-STAT5 (S726/731) in MV411 and MOLM-13 cells treated with the indicated concentrations of romaciclib and/or VEN for 24 hours. Representative western blots of two independent experiments are shown (except for 0.1 uM VEN concentration where n = 1).

Considering the mechanism of action of VEN,^5^ we next investigated whether this synergistic effect was associated with increased apoptosis. As expected, treatment with VEN alone induced apoptosis in both MV4-11 and MOLM-13 cells. Notably, co-treatment with romaciclib dose-dependently enhanced VEN-induced apoptosis in MV4-11 (after 96-hour exposure) and MOLM-13 cells (after 24-hour exposure) (**Figure 1C, Supplementary Figure S1A**).

To further explore the mechanistic basis of this synergy, we examined the level of relevant BCL-2 family proteins, particularly MCL-1 and Bcl-2 Interacting Mediator of cell death (BIM).

MCL-1 protein levels were reduced in both cell lines following VEN monotherapy, which was further enhanced by the RVU120+VEN combination (**Figure 1D**). The observed loss of MCL-1 coincided with increased cleavage of caspase-3, in line with the synergistic induction of apoptosis (**Figure 1D**). BIM was upregulated by romaciclib monotherapy, but this effect was not maintained in the combination treatment. Consistent with our previous findings,^20^ CDK8/19 inhibition with romaciclib led to a dose-dependent decrease of STAT5 phosphorylation at the S726/731 residues (**Figure 1D**). Moreover, total STAT5 levels were either undetectable or markedly reduced following RVU120+VEN treatment, leading to a complete loss of STAT5 phosphorylation (**Figure 1D**, **Supplemental Figure S1B**).

### Romaciclib and VEN combination induces caspase-dependent cleavage of MCL-1 and downregulates inflammatory signaling in VEN-sensitive AML cells

To identify molecular mechanisms underlying MCL-1 downregulation, we first examined the effect of romaciclib and/or VEN on MCL1 gene expression. RT-qPCR analysis revealed that neither the combination nor either agent alone markedly affected MCL1 transcript levels in MV4-11 and MOLM13 cells (**Supplemental Figure S2A**), suggesting a posttranscriptional mechanism.^22,23^ We observed that the RVU120+VEN combination reduced MCL-1 protein expression in parallel with a concurrent increase in S159 phosphorylation beyond VEN-inducible levels (**Figure 2A**). Further mechanistic studies revealed that the treatment with proteasome inhibitor MG-132 failed to rescue MCL-1 expression following RVU120+VEN treatment (**Supplemental Figure S2B**), excluding proteasomal degradation as a likely mechanism of MCL-1 downregulation in this context. Instead, given that the observed loss of MCL-1 was accompanied by increased caspase-3 activation (**Figure 2B**), we next investigated the role of caspases in this process. Consistent with previous reports on caspase-dependent degradation of MCL-1,^24–27^ we found that the pan-caspase inhibitor Z-VAD-FMK prevented RVU120+VEN-induced cleavage of MCL-1, restoring its protein levels in both MV4-11 and MOLM-13 cells (**Figure 2B**). Collectively, these findings demonstrate that in AML cells, RVU120+VEN combination promotes caspase-mediated and proteasome-independent MCL-1 degradation.

**Figure 2.**
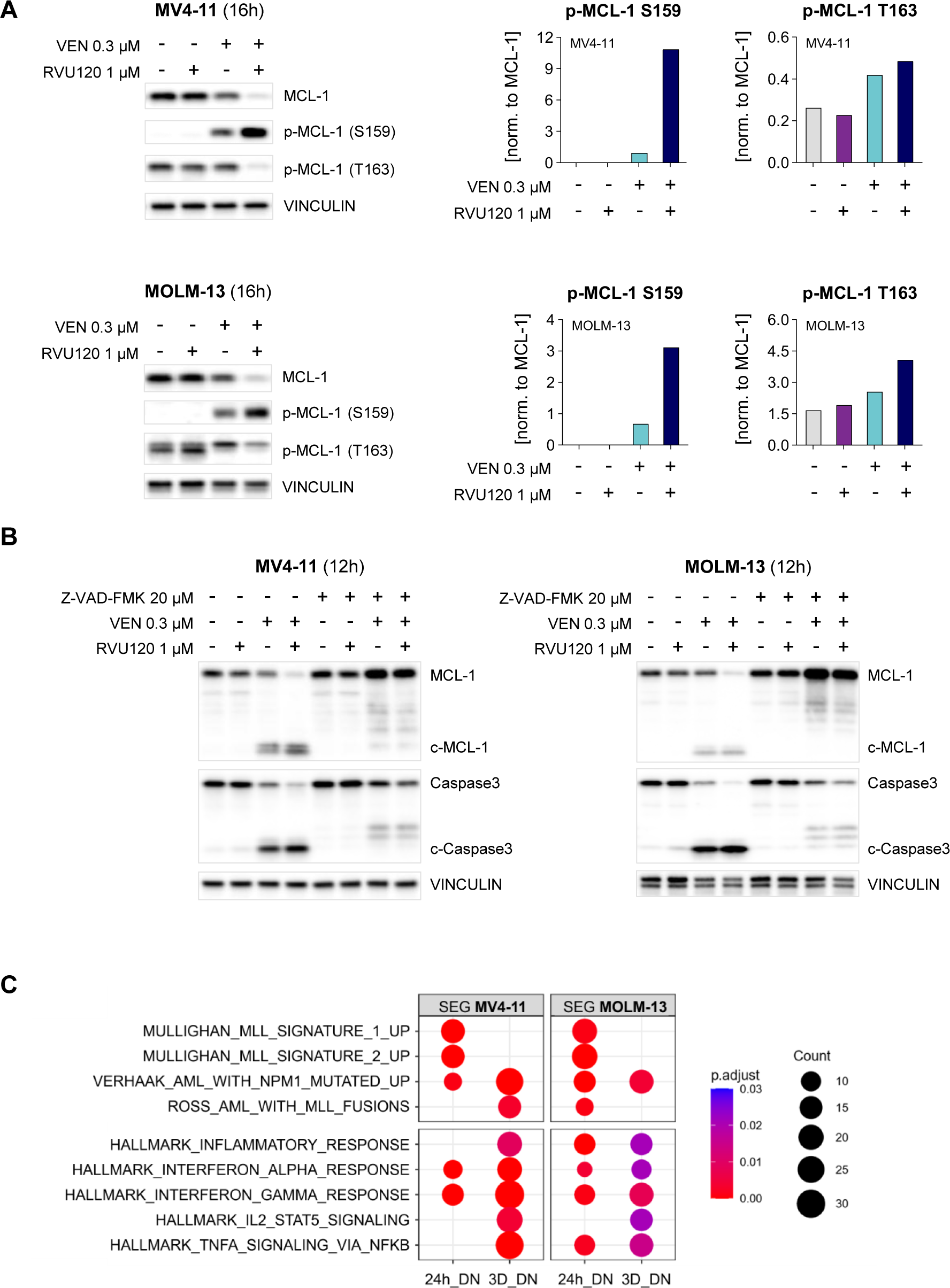
Combination of romaciclib and VEN induces caspase-dependent MCL-1 degradation in VEN-sensitive AML cells. **(A)** Protein expression of MCL-1, p-MCL-1 (S159), and p-MCL-1 (T163) in MV4-11 and MOLM13 cells treated with the indicated concentrations of romaciclib and/or VEN for 16 hours. The bar graphs show quantification of p-MCL-1 (S159) and p-MCL-1 (T163) levels. Representative western blots of two independent experiments are shown. **(B)** Protein expression of MCL-1 and cleaved MCL-1 (cMCL-1), caspase-3 and cleaved caspase-3 (c-caspase-3) in MV4-11 and MOLM-13 cells treated with the indicated concentrations of romaciclib and/or VEN for 16 hours, with or without pretreatment with pan-caspase inhibitor Z-VAD-FMK (30 min, 20 µM). Representative western blots of two independent experiments are shown. **(C)** Enrichment of the selected Hallmark/C2 collection gene sets within the genes synergistically downregulated by the combination of romaciclib (100 nM) + VEN (10 nM) in MV4-11 and MOLM-13 cells at 24 hours and 72 hours. SEG – synergistically expressed gene, DN – downregulation.

RNA-Seq profiling (**Supplemental Figure S3**) further revealed that romaciclib and VEN synergistically downregulated genes associated with inflammatory signaling such as the inflammatory response, IFN-a/g response, IL2-STAT5 signaling, and TNF-a signaling via NF-kB (**Figure 2C**). Considering that MV4-11 and MOLM-13 cells represent MLL-rearranged AML (MLL-AF4 and MLL-AF9 fusions, respectively), we found that the RVU120+VEN combination reversed oncogenic signaling signatures driven by MLL rearrangements in both cell lines (**Figure 2C**).

In summary, these results identify both post-translational and transcriptional mechanisms likely contributing to the antileukemic synergy between romaciclib and VEN in VEN-sensitive AML cells.

### Romaciclib and VEN combination is effective in primary AML patient samples ex vivo and in AML MV4-11 cell line xenograft

To evaluate the therapeutic potential of CDK8/19 and BCL-2 coinhibition, we first assessed ex vivo responses in a panel of PDCs from patients with de novo and R/R AML (n = 11). PDCs mutation status and treatment history are shown in **Supplemental Table S2**. To optimize the treatment regimen, we compared two dosing schemes: a continuous co-treatment with romaciclib and VEN for 96 hours vs a sequential co-treatment with VEN added only for the final 24 hours. In both conditions, monotherapies with romaciclib or VEN significantly decreased cell viability and induced apoptosis, though responses varied among PDCs ( **Figure 3A-B**).

**Figure 3.**
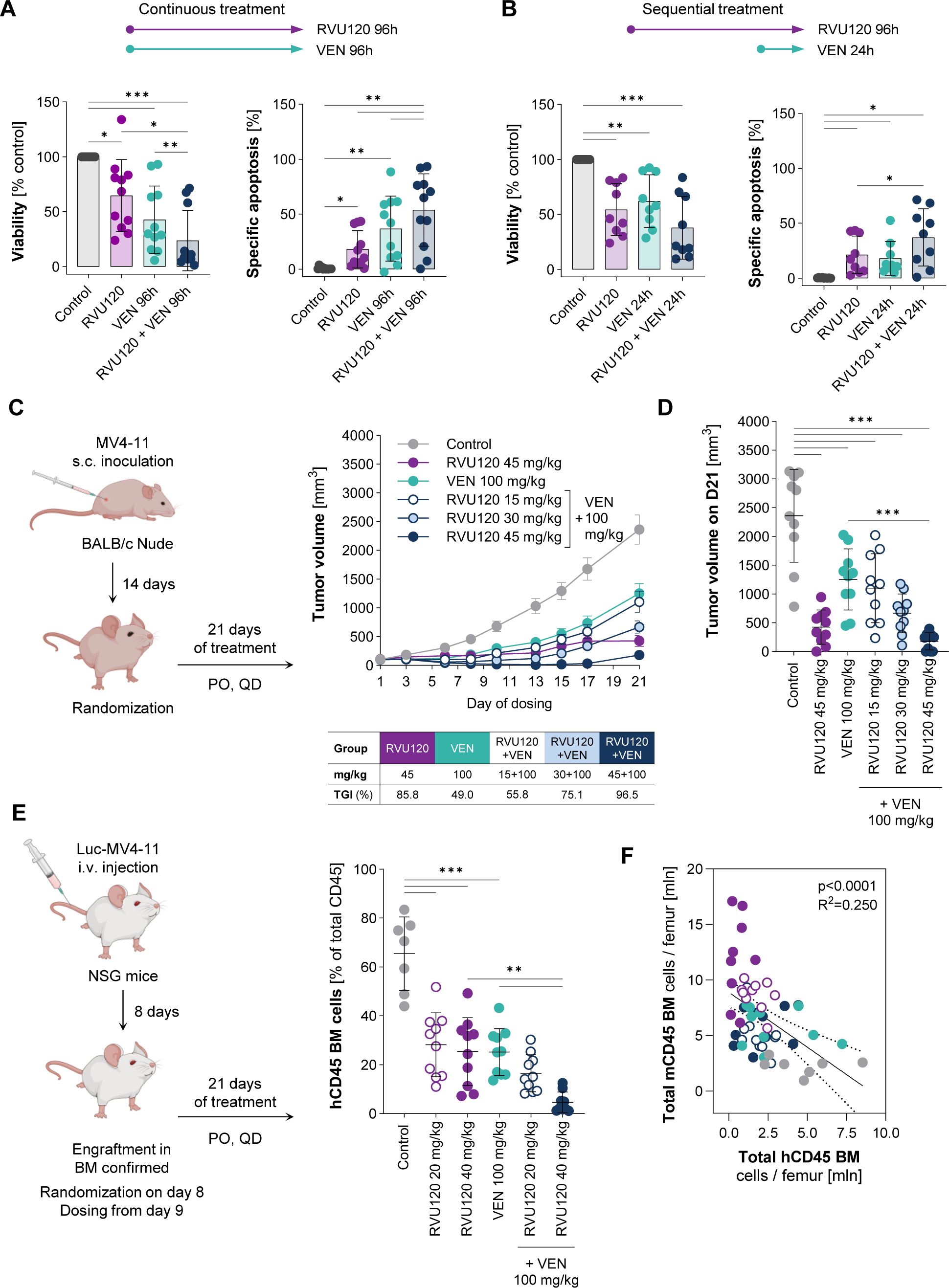
Romaciclib and VEN combination is effective in primary AML patient samples ex vivo and in AML MV4-11 xenograft models. **(A, B)** Cell viability and apoptosis induction in primary AML patient-derived cells treated (A) concomitantly with 1 μM romaciclib, 100 nM VEN, or their combination for 96 hours (n = 11), or (B) sequentially with 1 μM romaciclib for 72 hours, followed by 100 nM VEN for the final 24 hours (n = 9). *p < 0.05, **p < 0.01, ***p < 0.001 (RM one-way ANOVA with Tukey’s multiple comparisons test). **(C)** Schematic outline of the in vivo study with MV4-11 subcutaneous xenografts. Tumor volumes over time for the indicated treatment arms. Data are mean ± SEM. The TGI values are provided in the table. **(D)** Tumor volumes of individual mice after 21 days of treatment. ***p < 0.001 (one-way ANOVA with Tukey’s multiple comparisons test as indicated). **(E)** Schematic outline of the in vivo study with MV4-11 orthotopic xenografts. The percentage of human CD45+ leukemic cells in the bone marrow after 21 days of treatment. **p < 0.01, ***p < 0.001 (one-way ANOVA with Tukey’s multiple comparisons test as indicated). **(F)** Pearson correlation of the total murine (mCD45) and human (hCD45) CD45⁺ bone marrow (BM) cell numbers per femur. Each point represents an individual mouse, with colors indicating different treatment arms. The coefficient of determination (R²) and p-value are shown in the graph. PO – oral gavage, QD – daily administration

Combination of romaciclib and VEN enhanced the antileukemic effect of either monotherapy in 8 out of 11 PDC samples in the 96-hour continuous scheme (**Figure 3A**). A similar trend, although not statistically significant, was observed in the sequential co-treatment (**Figure 3B**). To determine whether the in vitro efficacy of the RVU120+VEN combination translated into in vivo models, we first utilized a subcutaneous MV4-11 xenograft model (**Figure 3C**). VEN alone inhibited tumor growth by 49% (that is below the effective threshold of 58%), while monotherapy with romaciclib at 45 mg/kg was followed by 86% tumor growth inhibition (TGI). Co-administration of romaciclib enhanced the anti-tumor activity of VEN in a dose-dependent manner, with the top dose showing 97% TGI. Complete remissions were noted in animals dosed with romaciclib at 45 mg/kg (one mouse) and in animals receiving a combined treatment with VEN and romaciclib at the top dose (three mice) (**Figure 3C-D**).

To corroborate these findings, we also assessed the efficacy of the RVU120+VEN combination in an orthotopic Luc-MV4-11 cell line xenograft model. As shown in **Figure 3E**, monotherapies with romaciclib (at both 20 mg/kg and 40 mg/kg) or VEN (at 100 mg/kg) significantly reduced the percentage of hCD45+ leukemic cells in BM compared to the control group. Notably, the RVU120+VEN combination further reduced the fraction of hCD45+ cells compared to either monotherapy. This resulted in the lowest total number of hCD45+ leukemic cells/femur in the romaciclib (40 mg/kg) +VEN treatment arm and was paralleled by the host BM recovery (**Figure 3E-F**, **Supplemental Figure S4A-B**).

In summary, we demonstrated that the RVU120+VEN combination exhibits synergistic antileukemic activity across multiple AML models, including cell lines, PDCs, and in vivo AML xenografts.

### Romaciclib and VEN demonstrates synergistic antileukemic activity in VEN-resistant AML cells

Next, we assessed the potential of the RVU120+VEN combination in cell-intrinsic and cell-extrinsic models of VEN-resistant AML. Considering that romaciclib inhibits STAT signaling (**Figure 2B-C** and our previous findings^20^) and that JAK-STAT-dependent stromal factors reduce AML sensitivity to VEN, we mimicked the protective effect of BM microenvironment using HS-5 stromal cell-conditioned medium (CM).^28^ In contrast to romaciclib, VEN activity was markedly diminished in AML cell lines in the presence of HS-5 CM (**Figure 4A**). RVU120+VEN treatment showed high synergy and superior efficacy under these conditions (**Figure 4B-C**), suggesting that the combination can overcome the inhibitory effect of BM stroma.

**Figure 4.**
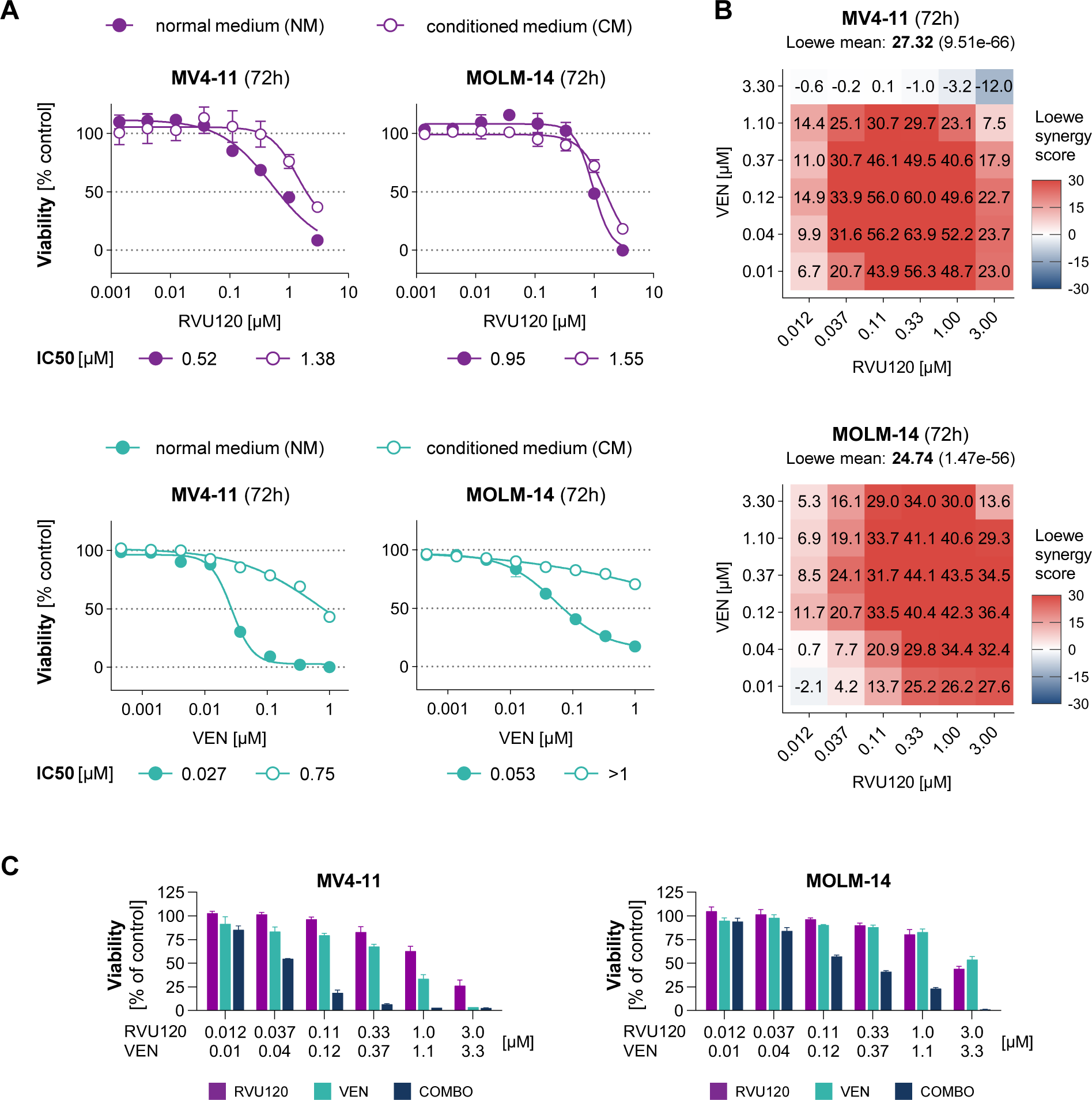
Romaciclib in combination with VEN overcomes stroma mediated VEN-resistance. **(A)** Dose-response viability assay for MV4-11 and MOLM-14 cells treated with increasing concentrations of romaciclib or VEN for 3 days, in the absence or presence of HS-5 stromal cell-conditioned medium. Data are mean ± SD of technical triplicates from a representative experiment (n = 3). **(B)** Loewe synergy scores calculated from the effect of RVU120+VEN combination on viability of MV4-11 and MOLM-14 cells treated with the indicated concentrations of drugs for 3 days, in the presence of HS-5 stromal cell-conditioned medium. Loewe scores of > 0 are regarded as likely synergistic. Data from technical triplicates from a representative experiment are shown (n = 2). **(C)** Bar graph showing matched cell viability at select doses. Data are mean ± SD of technical triplicates from a representative experiment (n = 2).

A recent integrative meta-analysis of AML patients from the BeatAML1 study identified four transcriptionally distinct clusters of VEN resistance (VR_C1-4).^14^ To assess the efficacy of RVU120+VEN in this context, we tested the combination in a panel of nine VEN-resistant AML cell lines, representing three of these resistance clusters VR_C1-3 (VR_C4 is transcriptionally similar to VEN sensitive cluster) (**Figure 5A**).^14^ The RVU120+VEN combination exhibited a synergistic antileukemic effect in AML cell lines from the VR_C1 and VR_C2 clusters (**Figure 5B-C, Supplemental Figure S5**). The strongest synergy was observed in VR_C2 cell lines (mean Loewe synergy scores > 10 in 100% of cell lines) whereas no synergy was found in VR_C3 representatives. Notably, VR_C1/2 cell lines exhibiting synergy predominantly represented a monocytic disease phenotype, according to French-American-British (FAB) classification (**Figure 5A**).

**Figure 5.**
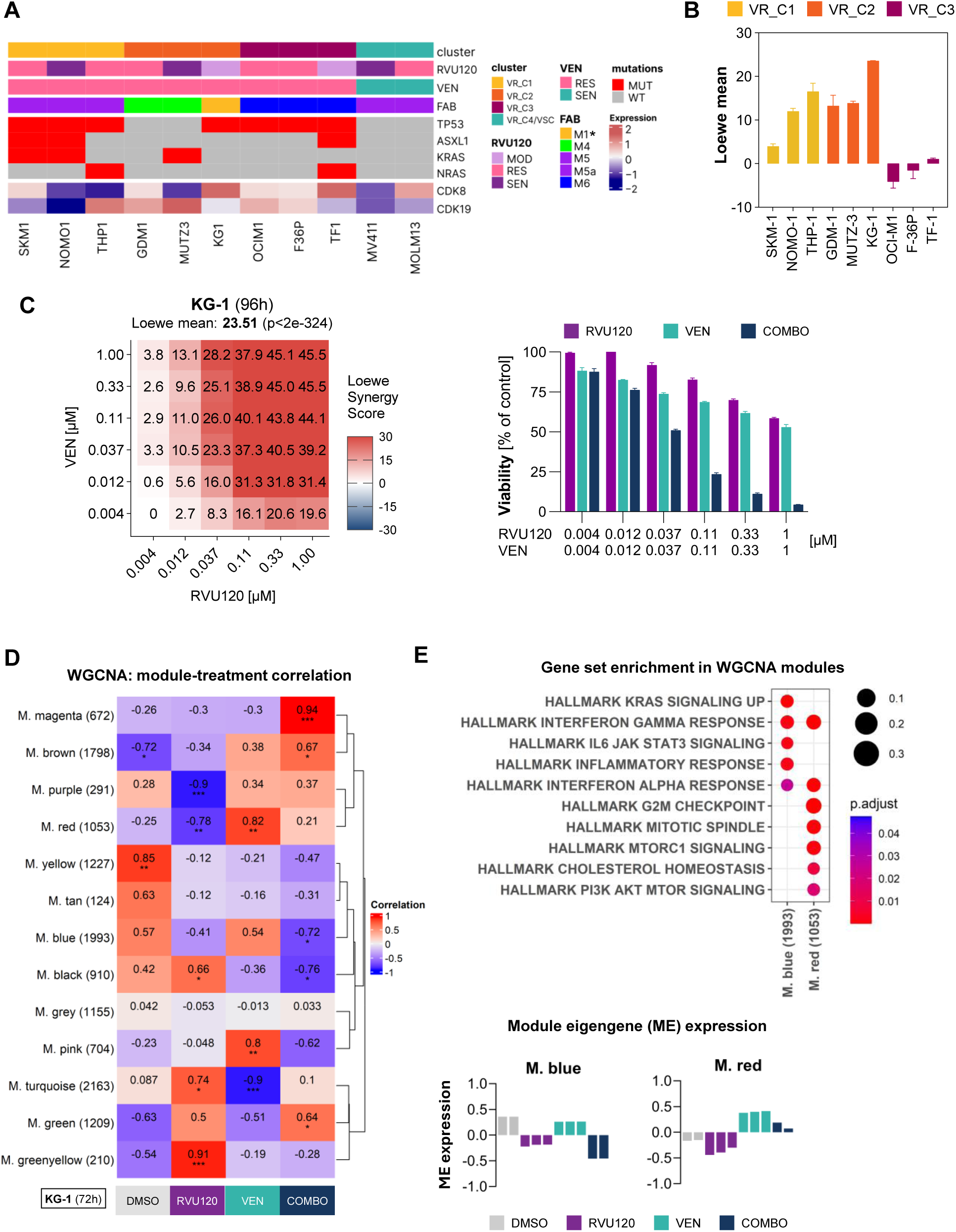
Romaciclib and VEN demonstrates synergistic antileukemic activity in VEN-resistant AML cells. **(A)** Characteristics of clustered AML cell lines. The table presents the sensitivity of AML cell lines to romaciclib and VEN monotherapy, along with their FAB classification, mutation status, relative transcript levels of CDK8 and CDK19. Abbreviations: SEN = sensitive, RES = resistant, MUT = mutated, WT = wild type. *Classification based on Skopek et al., Int J Mol Sci., 2023^29^ **(B)** Mean Loewe synergy scores for the RVU120+VEN combination in VEN-resistant AML cell lines. Loewe scores of > 0 are regarded as likely synergistic. Data are mean ± SD of two independent experiments. **(C)** Loewe synergy scores calculated from the effect of the RVU120+VEN combination on viability of KG-1 cells treated with the indicated romaciclib/VEN concentrations for 96 hours. Loewe scores of > 0 are regarded as likely synergistic. The bar graph shows matched cell viability at select doses. Data from triplicates from a representative experiment are presented (n = 2). **(D)** Heatmap showing correlations between the treatment groups and modules of co-expressed genes identified by a network-based analysis (WGCNA) of RNA-Seq data from KG-1 cells treated with 100 nM romaciclib, 10 nM VEN or their combination for 72 hours. **(E)** Enrichment of the select Hallmark collection gene sets in the indicated WGCNA modules (top). Bar plots showing module eigengene (ME) expression across the treatment groups (bottom).

To elucidate the molecular mechanisms underlying this synergistic effect, we performed RNA-Seq profiling in KG-1 cells^29^ —a well-characterized model with documented in vivo efficacy and pharmacology data for romaciclib.^20^ Network-based analysis of the RNA-Seq data revealed distinct modules of coexpressed genes that correlated with either monotherapy or RVU120+VEN co-treatment (**Figure 5D**). Specifically, we focused on gene sets associated with VEN resistance found in two modules: (a) the red module (M. red), which includes genes induced by VEN but suppressed by romaciclib and the RVU120+VEN combination; and (b) the blue module (M. blue), which contains genes downregulated by the RVU120+VEN combination. These modules were significantly enriched for several inflammatory pathways including IL6-JAK-STAT3 signaling, IFN-a/g response, and inflammatory signaling and metabolic pathways such as KRAS signaling and PI3K-AKT-MTOR (**Figure 5E**).

### Romaciclib and VEN synergize ex vivo in VEN resistant PDX model via JAK/STAT3 and PI3K-AKT pathways

To confirm this mechanism of synergy in primary setting, we examined the activity of RVU120+VEN combination in a VEN-resistant PDX model. The PDX model AMLX12 was classified based on its transcriptional profile into the VR_C2 cluster of resistance (**Supplemental Figure S6**). AMLX12 contained different immunophenotypically-defined subpopulations: an immature, potentially enriched in CD34+CD38- and CD123+(CD34+CD38+) LSC-like cells, and a more mature, CD45RA+CD11b+ cells (**Figure 6A**). AMLX12 exhibited primary resistance to VEN monotherapy (consistent with its cluster assignment) and was insensitive to romaciclib alone (**Figure 6B**). Despite this, the RVU120+VEN combination demonstrated functional synergy in AMLX12, dose-dependently reducing cell viability after 72 hours of treatment (**Figure 6B**).

**Figure 6.**
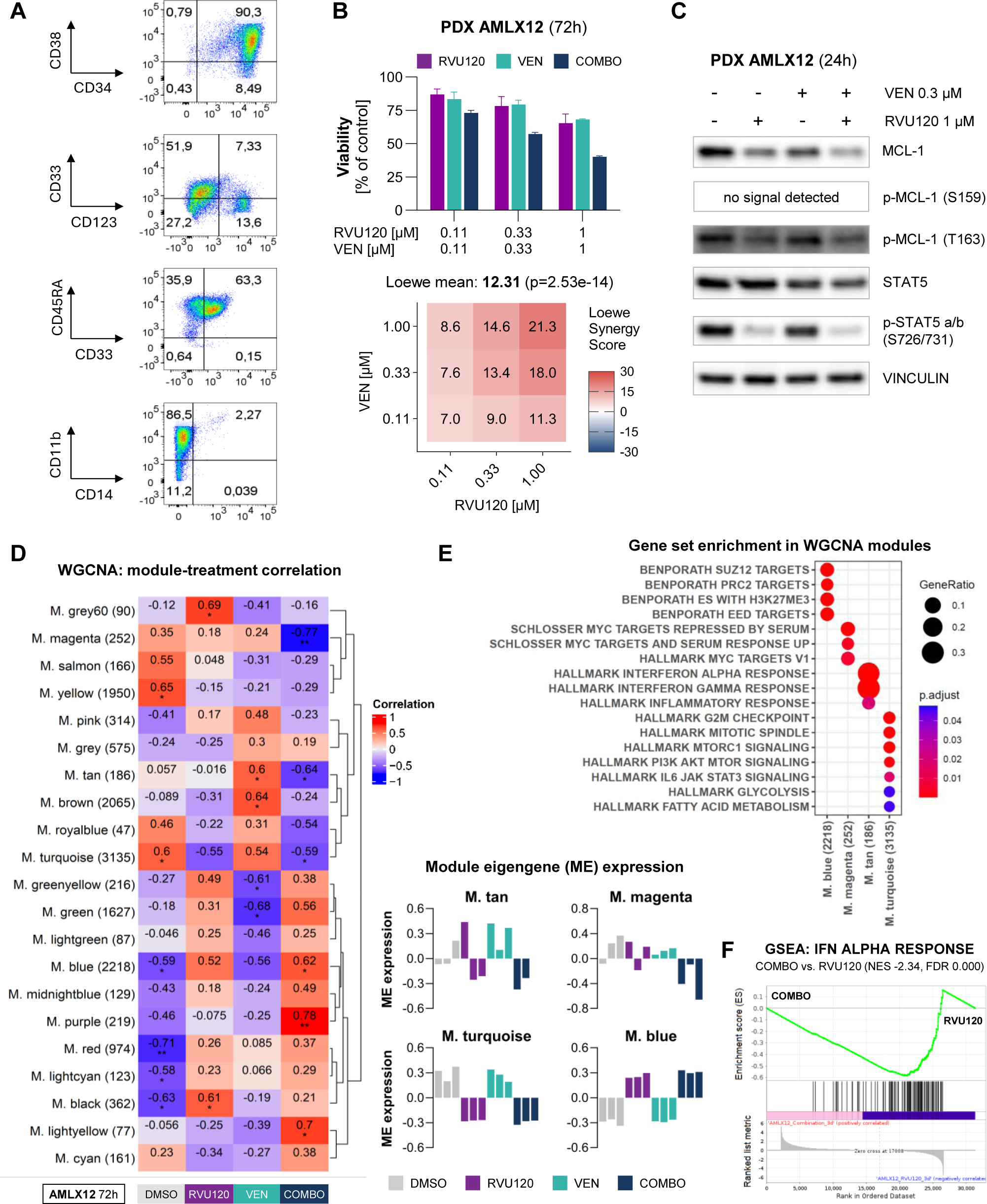
Romaciclib and VEN demonstrates synergistic antileukemic activity in a primary PDX model of VEN-resistant AML. **(A)** Immunophenotypic profile of PDX-AMLX12. The plots show the expression of surface markers: CD34, CD38, CD33, CD123, CD45RA, CD11b, and CD14, in the hCD45+ cell population. Representative plots of two independent experiments are shown. **(B)** Loewe synergy scores calculated from the effect of RVU120+VEN combination on viability of PDX-AMLX12 cells treated with the indicated romaciclib /VEN concentrations for 72 hours. Loewe scores of > 0 are regarded as likely synergistic. The bar graph shows matched cell viability. Data from a technical triplicate are presented (n = 1). **(C)** Protein expression of MCL-1, p-MCL-1 (S159), p-MCL-1 (T163), STAT5 total and p-STAT5 (S726/731) in PDX-AMLX12 cells exposed to the indicated concentrations of romaciclib and/or VEN for 24 hours (n = 1). **(D)** Heatmap showing correlations between the treatment groups and modules of co-expressed genes identified by a network-based analysis (WGCNA) of RNA-Seq data from PDX-AMLX12 cells treated with 100 nM romaciclib, 10 nM VEN, or their combination for 72 hours. **(E)** Enrichment of the selected Hallmark and C2 collections gene sets in the indicated WGCNA modules (top). Bar plots showing module eigengene (ME) expression across the treatment groups (bottom).**(F)** GSEA running plot showing negative enrichment of the IFN-a response Hallmark gene set in COMBO (RVU120+VEN)-treated PDX-AMLX12 cells compared to romaciclib monotherapy.

Consistent with our observations in cell lines, this antileukemic effect in AMLX12 was accompanied by MCL-1 downregulation and reduced S726/731 STAT5 phosphorylation (**Figure 6C**). These findings suggest that, at least in part, common underlying mechanism may drive RVU120+VEN synergy across different AML models.

Indeed, RNA-Seq analysis of PDX-AMLX12 responses to romaciclib and/or VEN identified a distinct downregulation of inflammatory pathways: IFN-a/g and inflammatory response shown in the tan module (M. tan), mirroring transcriptional changes observed in KG-1 cells (**Figure 5E**). This anti-inflammatory effect was corroborated by an independent gene set enrichment analysis comparing the RVU120+VEN combination vs romaciclib monotherapy (**Figure 6F**). Additionally, pathways implicated in VEN resistance were downregulated by the combination treatment, shown in the turquoise module (M. turquoise). These included IL6/JAK-STAT3 signaling, metabolism-associated pathways (glycolysis, mTORC1 signaling, PI3K-AKT signaling), and cell cycle-related pathways (G2M checkpoint, mitotic spindle) ^14,30^ (**Figure 6D-E**). Notably, some of these pathways were already downregulated by romaciclib monotherapy. Furthermore, romaciclib treatment reversed Polycomb Repressive Complex 2 (PRC2)-mediated repression associated with a stem cell-like phenotype. This is evidenced by the enrichment of the PRC2 gene set and its components such as Suppressor of Zeste 12 (SUZ12), Embryonic Ectoderm Development (EED), embryonic stem cell (ES) shown in the blue module (**Figure 6D-E**).

Together, these findings from the PDX-AMLX12 model provide mechanistic proof-of-concept validation for the antileukemic synergy of romaciclib and VEN in a primary model of VEN-resistant AML.

### Combination of romaciclib and VEN suppresses leukemia burden in vivo in VEN-resistant PDX model

Finally, to investigate the clinical relevance of these findings, we tested the efficacy of romaciclib and VEN in vivo using VEN-resistant PDX model (VR-PDX-Sample-1, PDX-S1). This model was developed from primary PBMC of a MDS patient that further progressed to AML and has previously been used in myeloid leukemia studies.^31^ Similar to AMLX-12, PDX-S1 transcriptionally represent VR_C2 cluster of VEN resistance (**Supplementary Figure 6**). PDX cells were engrafted into NSG mice, with successful engraftment confirmed 5 weeks post-injection. Mice were then treated according to the scheme provided in **Figure 7A** and analyzed using gating strategy shown in **Figure 7B**. As anticipated, PDX-S1 model did not respond to VEN monotherapy (**Figure 7C),** whereas romaciclib alone decreased the percentage of total human and myeloid PDX cells compared to VEN. Importantly, the RVU120+VEN combination overcame VEN resistance and achieved the greatest reduction in hCD33+ myeloid cells compared with either monotherapy (**Figure 7C**). Leukemia initiating cells (LIC) are often cause of relapses, consequently, targeting this population could improve patient outcomes. We therefore characterized AML cells to define a LIC-enriched fraction (**Figure 7B**). Combination treatment significantly lowered the frequency of LIC population in the BM, as indicated by decrease in the percentage of hCD45+Lin-CD34+ cells (**Figure 7D**). Collectively, these in vivo findings strongly suggest that romaciclib combined with VEN may represent an effective therapeutic strategy for patients with VEN-resistant AML.

**Figure 7.**
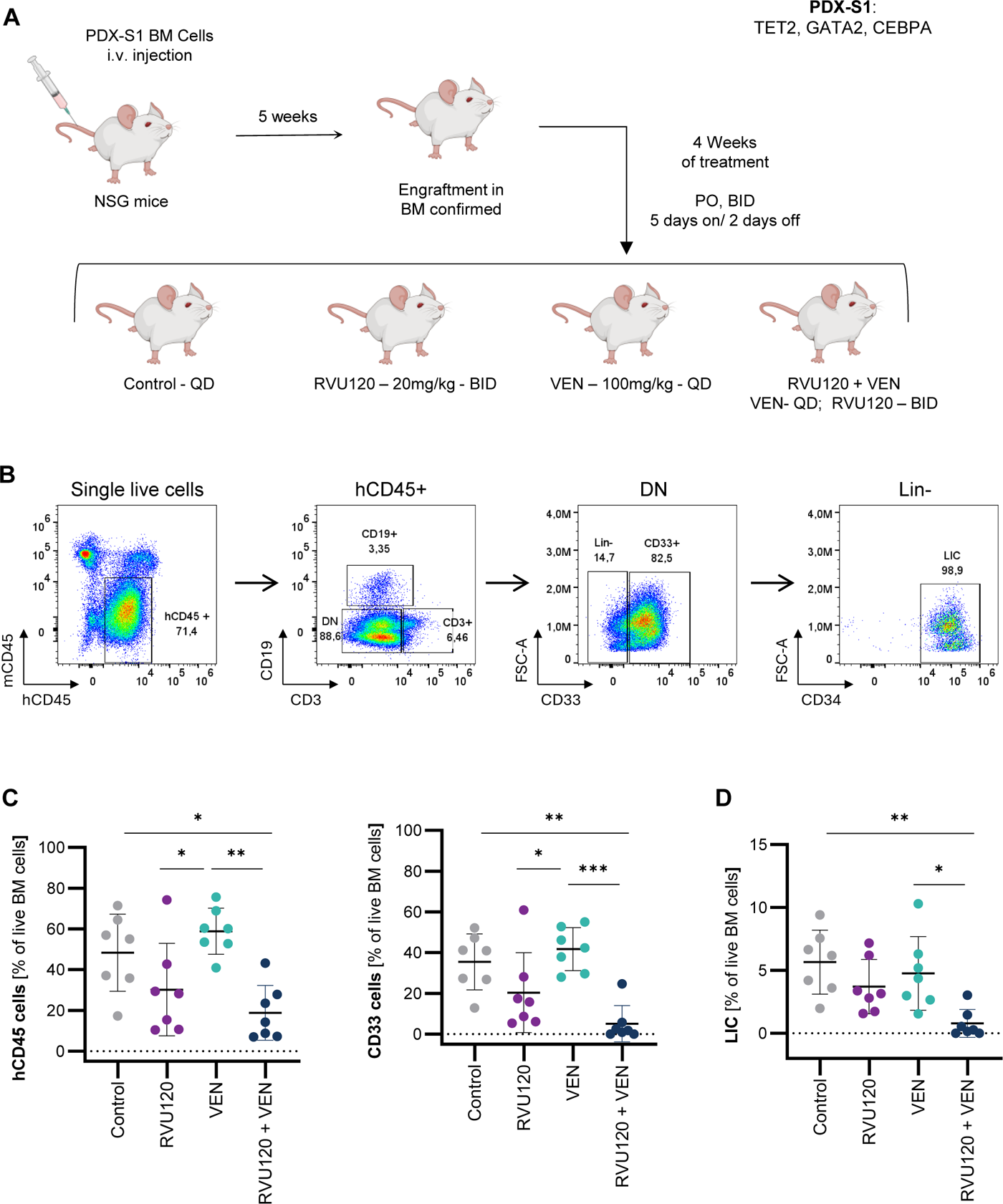
Romaciclib and VEN combination suppresses leukemia burden in vivo in VEN-resistant PDX model. **(A)** Schematic illustrating the transplantation strategy used to establish the in vivo VEN-resistant PDX-S1 model (TET2, GATA2, CEBPA), along with the treatment groups in NSG mice. **(B)** Representative flow cytometry plots depicting the gating strategy for analysis of mCD45-hCD45+ cells, hCD45+hCD33+ cells and hCD45+lin-CD34+ cells following 4 weeks of treatment. **(C-D)** Scatter plot showing the percentages of live hCD45+ cells and hCD33+ cells **(C)** and CD34+ Cells **(D)** in BM from PDX-S1 model under different treatment conditions (Mean ± SD, n=7; Ordinary one-way ANOVA with post hoc Tukey correction). *p < 0.05, **p < 0.01, ***p < 0.001

## Discussion

Despite the recent progress, the majority of patients experience refractory or relapse disease upon VEN and hypomethylating agent treatment. This highlights the ongoing need to refine treatment strategies to improve response rates and prolong AML remission.

In this study, we demonstrated robust synergistic anti-leukemic activity of the RVU120+VEN combination across multiple AML models. This effect was validated in vitro in AML cell lines, ex vivo in primary patient-derived samples and in vivo using VEN-sensitive and -resistant models.

Mechanistically, RVU120+VEN treatment resulted in caspase-dependent cleavage of MCL-1, independent of proteasomal degradation. MCL-1 is a key mediator of suboptimal VEN efficacy and resistance.^6,32^ The combination greatly increased MCL-1 phosphorylation at serine 159, which is known not only to be involved in proteasomal degradation but also required for caspase-dependent cleavage.^25,33,34^ Importantly, cleaved MCL-1 fragments also possess pro-apoptotic properties, amplifying cell death.^24–27^ In addition, the combination decreased total STAT5 levels. Given that STAT5 regulates MCL-1 expression at the transcriptomic level and also via PI3K pathway, particularly in FLT3-ITD AML, decrease of STAT5 may further contribute to the observed synergy.^35^ Although our experiments in FLT3-ITD cell lines did not show significant reductions in MCL-1 mRNA, the potential involvement of STAT5/PI3K-mediated regulation of MCL-1 warrants further investigation. Finally, transcriptomic analysis in VEN sensitive setting revealed that RVU120+VEN synergistically downregulated inflammatory pathways including IL2/STAT5 and TNF/NF-κβ. TNF/NF-κβ pathway is involved in resistance to VEN treatment in T-cell acute lymphoblastic leukemia and AML models and inhibition of this pathway synergizes with BCL-2 inhibition.^36,37^ Inflammatory pathways such as TLR/MyD88/NF-κB are also upregulated in MLL-rearranged leukemia, and together with MLL fusions phenotype, are associated with stem-like traits, self-renewal, and poor prognosis.^38–41^ In addition to inflammatory signaling, romaciclib with VEN synergistically repressed MLL fusion-driven oncogenic programs, suggesting the potential to reverse the stem cell-like phenotype, particularly in MLL-AF4/AF9 cells.

We also addressed the challenge of microenvironment-driven resistance. AML cell lines sensitive to VEN in vitro often lose responsiveness in vivo due to the BM niche.^42,43^ Further, stromal cells modulate BCL-2 family protein expression, particularly BCL-2 and BCL-xl.^17,28^ Interestingly, MCL-1 contributes to drug resistance beyond its anti-apoptotic function by regulating cellular metabolism and promoting stroma-AML interactions via the CXCR4 and CD44 axis.^44^ The CXCR4 axis also enables leukemic cells to outcompete healthy hematopoietic stem cells for niche occupancy.^45^ Consequently, MCL-1 downregulation caused by RVU120+VEN may enhance therapeutic outcomes through multiple mechanisms beyond apoptosis activation.^17,28,40^ In our study, using the HS-5 stroma model, VEN activity was significantly reduced, whereas romaciclib retained efficacy. The combination was highly synergistic and overcame stroma-mediated resistance to VEN. Previous studies have shown that HS-5 stromal cells secrete cytokines and growth factors, such as GM-CSF and G-CSF, which have been shown to confer resistance to VEN through activation of JAK/STAT signaling, including downstream STAT5.^28^ This mechanism may underlie the synergy observed with romaciclib and VEN, as the combination downregulated IL6/JAK/STAT signaling and markedly reduced total STAT5 protein levels.

Recent studies highlight the need for integrated approaches that extend beyond genomic profiling to fully elucidate the mechanism of therapy resistance, particularly in R/R AML.^13,46^ In this context, the combination of romaciclib and VEN demonstrated strong synergy in two (VR_C1 and VR_C2) of the three tested VEN-resistant transcriptional clusters identified by Mohanty et al.^14^ Moreover, AML cell lines in which synergy was observed belong to the M4 and M5 subtypes of the French–American–British (FAB) classification. In contrast, no synergy was observed in VR_C3, enriched for the erythroleukemia/M6 subtype. These findings are consistent with previous reports demonstrating reduced sensitivity of monocytic/mature AML blasts to VEN-based regimens.^10,46,47^. Furthermore, the combination therapy effectively overcame VEN resistance in the PDX-S1 model with a primitive (CD34⁺) phenotype, as evidenced by a marked reduction in both blast and LIC populations. Thus, RVU120+VEN may be active in AML subsets characterized by both monocytic-like and immature/primitive blasts. Mechanistically, RVU120+VEN suppressed several resistance-associated pathways, including IL6/JAK/STAT3, PI3K/AKT/MTOR, and inflammatory signaling — previously identified as key mediators of VEN resistance.^14,30^ Importantly, romaciclib monotherapy was sufficient to downregulate inflammatory gene signatures, particularly interferon alpha (IFNα) and gamma (IFNγ). IFNγ signaling is associated with the AML maturation state, showing the highest activity in monocytic-like leukemic blasts and VEN resistance.^41^

Suppression of the IL6/JAK/STAT3 pathway is consistent with the established role of CDK8 as a regulator of STAT3 expression and its phosphorylation at serine 727.^48^ Notably, both total STAT3 and its phosphorylated form (S727), linked to mitochondrial dysfunction, are upregulated in VEN-resistant cells along with its downstream target – MCL-1.^49^ Collectively, these findings suggest that CDK8 inhibition by romaciclib may also overcome resistance through the STAT3/MCL-1 axis while simultaneously restoring mitochondrial function.

In summary, our findings provide, for the first time, a comprehensive understanding of the mechanism of action of RVU120+VEN both in sensitive and VEN-resistant AML models, supporting its potential as a promising therapeutic strategy across genetically and transcriptionally diverse AML subtypes. Importantly, RVU120+VEN combination may offer a valuable treatment option for patients with relapse/refractory disease who otherwise lack effective alternatives.

## Acknowledgments

The authors thank Jessica T. Swann, a senior scientific editor at MD Anderson Cancer Center, for providing editorial assistance. This research was supported in part by grant from the Polish Medical Research Agency (ABM) 2022/ABM/06/00002 and the National Centre for Research and Development (NCBR) POIR/01.02.00-12-32/15 (Ryvu Therapeutics). This work was supported, in part, by grants from the Cancer Prevention and Research Institute of Texas (CPRIT) and the National Cancer Institute (NCI) at National Institutes of Health (NIH) R01CA231364 and Leukemia SPORE P50 CA100632 (N.B.). This research was supported by the National Science Centre, Poland (2021/43/B/NZ5/03345) (N.B.) and 2018/29/B/NZ5/01471 (P.J). PDX-AMLX12 development was supported by the National Science Center grant (2018/30/E/NZ2/00801) (M. Mi.).

## Authorship Contributions

U.P., T.R., and M.Ma. were responsible for conceptualization, methodology, investigation; N.B. provided critical guidance on study design; U.P., M.O., J.W., K.W., S. C., M.M, V.M., D.C., E.A., A.G., A.M., H.Z., M.C., E.K., Z. S., and N.B. performed experiments and/or analyzed the data; M.K., M.Mi., P. J., and A.S. provided resources and/or guidance on experimental design; U.P., M.O., J.W., and T.R. prepared the original manuscript draft; U.P., M.Ma., and N.B. reviewed and edited the manuscript; T.R., and M.Ma. supervised the study; T.R., and M.Ma. were responsible for project administration; and all authors have read and agreed to the published version of the manuscript.

## Conflict-of-interest disclosures

U.P., M.O., J.W, K.W., M.M., D.C., E.A., A.G., A.M., P.J., T.R. and M.Ma. are current or former Ryvu Therapeutics employees and/or shareholders. In the past 2 years A. S. has received honoraria for educational activities from PeerView, ACHL, Great Debates, HemeOnc Pulse and Geron; advisory board fees from Kymera Therapeutics, Geron, Ryvu Therapeutics; research funding from Kymera Therapeutics and Ryvu Therapeutics and is Data Safety Monitoring Board Chair for Servier Pharmaceuticals. The remaining authors declare no competing financial interests.

## Notes

### Summary of Updates

Correct the misspelling of one of the co-authors name: Zuzanna Sandowska-Markiewicz and correct figure 7A: replace ROMA abbreviation with RVU120

