## Supplemental data for "Romaciclib, a CDK8/CDK19 inhibitor, can overcome venetoclax resistance through a combinatorial strategy"

### Supplemental Methods

#### Real-time RT-PCR

Total RNA was isolated using a NucleoSpin RNA Mini kit (Macherey-Nagel 740955), and cDNA was synthesized from 1 mg total RNA using a High-Capacity cDNA Reverse Transcription Kit (Thermo Scientific 4368813). Quantification of MCL-1 transcripts was done using Taqman probes (Life Technologies: MCL-1 Hs_01050896, TBP Hs_00427620) and a QuantStudio 6 instrument (Thermo Scientific). Relative gene expression was calculated using the comparative Ct method.

#### Subcutaneous MV4-11 xenograft model

The study was performed according to the protocol approved by the Institutional Animal Care and Use Committee (IACUC) of Shanghai ChemPartner following the guidance of the Association for Assessment and Accreditation of Laboratory Animal Care. After a minimum of 5 days acclimatization, mice were marked by ear coding and injected subcutaneously with 5 mln of MV4-11 (ATCC) viable cells suspended in 0.2 mL of serum-free Iscove's Modified Dulbecco's Mediumand Matrigel (1:1). The day of inoculation was denoted as day 1 of the study. Body weight (BW) and tumor volume (TV) measurements (in two dimensions using a caliper) were done three times per week. The results (not blinded) were distributed via a preformatted Excel file and a built-in GraphPad Prism. After 14 days of tumor formation, mice were randomly distributed into six experimental groups using a stratified randomization method, with an average TV of about 100 $mm^3$. Romaciclib was prepared in sterile water and dosed at 10 mL/kg (monotherapy) or 5 mL/kg (combined treatment) as a solution, and VEN was suspended in sterile water with 5% DMSO, 50% PEG300, and 5% Tween 80, and dosed at 10 mL/kg. Both drugs were administered once daily (QD) by oral gavage (PO) for three weeks. In the case of a combined dosing, romaciclib was administered 1-2 hours earlier than VEN. Animals were checked daily for morbidity and mortality. Any mouse reaching TV > 2000 mm3 or body weight loss (BWL) > 20% was euthanized; mice showing c.a. 10% BWL were half-dosed until recovery. The study was terminated after the last BW and TV measurements on day 21 of dosing.

#### MV4-11 orthotopic xenograft model

Animal welfare complied with the UK Animal Scientific Procedures Act 1986 in line with Directive 2010/63/EU. Anesthesia, husbandry, data management, and reporting were conducted in accordance with Crown Bioscience UK (CBUK) guidelines and standard operating procedures. After a minimum of 5 days acclimatization, mice were tagged with the use of transponder chips and injected intravenously via tail vein with 5 mln of MV4-11 (ATCC) viable cells, Luc-tagged at CBUK, suspended in 0.1 mL of PBS. The day of inoculation was denoted as day 0, and in vivo bioluminescence imaging (BLI; Spectrum CT imaging chamber) was used to indicate disease burden. Ventral and dorsal signals captured and analyzed using Living Image 4.3.1. software (Caliper LS, US). Based on bioluminescence-detectable engraftment on day 8, mice were randomized into six experimental groups using the matched distribution method, resulting in the average BLI of $1.8x10^7$ total flux (p/s). Sterile water with 5% DMSO, 30% PEG400, 0.5% Kolliphor EL, and 5% propylene glycol was used as a vehicle. Drugs were administered once daily (QD) by oral gavage (PO) with a 10 mL/kg per dose. Romaciclib was dosed as a solution, VEN was administered as a homogenous suspension, while romaciclib plus VEN were mixed 3:1 (v/v) and given to the animals as a homogenous suspension. Animal body weight and welfare were monitored at least once daily for the whole study duration. Data captured using StudyDirector (StudyLog System Inc.; not blinded) were exported to Microsoft Excel and GraphPad Prism for further analysis. Dosing holidays were applied to individual animals showing BWL > 10% and resumed after recovery to < 5% BWL. The humane endpoint was set at the level of BWL > 20%. The leukemic burden was monitored by BLI once a week, with the last measurement planned on day 21 of the dosing period. By this time, none of the animals had a total flux above the humane endpoint threshold (> $1x10^{10}$ p/s). After 21-22 days of treatment, 3-4 mice per day were sacrificed over three consecutive days to collect the material.

BM cells were isolated from the femurs of all animals in aseptic condition by flushing out BM cells with DPBS. After centrifugation and cell counting (automated cell counter), BM samples were suspended in the freshly prepared freeze media (10% DMSO in FBS) and immediately frozen at –80 °C. About $150x10^3$ cells from each vial were taken for flow cytometry analysis for human and murine CD45 expression (BioLegend: hCD45 APC 368512, mCD45 PE-Cy7 103114; Life technologies: Live/Dead fixable near-IR stain). The percentage of hCD45+ cells was calculated from the sum of hCD45+ and mCD45+ cells.

**AML patient derived xenograft study**

The study was performed under protocols approved by the institutional animal facility at Albert Einstein College of Medicine. Mice were monitored daily. 5 weeks post-transplantation mice were randomized into 4 treatment groups and dosed by oral gavage using a 5 days on / 2 days off schedule for 4 weeks:

- Control – Vehicle only (Formulation = 5% DMSO + 30% PEG400 + 0.5% Kolliphor EL + 5% propylene glycol + 59.5% water) administered PO daily.
- romaciclib – 20mg/kg – administered PO BID
- VEN (Selleck Chemicals, Cat #S8048) – 100mg/kg – administered PO once daily
- romaciclib (20mg/kg) + VEN (100mg/kg) – Co-administered following the same PO regimen

Control mice received vehicle formulation in the morning and water in the evening. Romaciclib group received 20mg/kg drug resuspended in formulation in the morning and 20mg/kg resuspended in water in the evening. VEN group received 100mg/kg VEN in the formulation in the morning and water in the evening. RVU120 + VEN group received 20mg/kg romaciclib along with 100mg/kg VEN in the formulation in morning and 20mg/kg romaciclib in water in the evening.

**RNA-Seq processing and data analysis**

Data were processed and analyzed with a custom version of the Nextflow pipeline based on nf-core/rnaseq v.3.14.0 (https://doi.org/10.5281/zenodo.1400710) of the nf-core collection of workflows[1], using software environments from the Bioconda[2] and Biocontainers[3] projects. In short, FASTQ files were aligned with STAR[4] to the GRCh38 reference genome using Gencode v.43[5] and quantified with salmon. If needed, mouse reads were removed using xengsort.[6] The pipeline was executed with Nextflow.[7] After data processing, 69 from 70 samples (outlier: AML_47) were eligible for further analysis. The raw RNA sequencing datasets and count tables generated in this study can be found in the GEO repository under accession code GSE292119.

**Differential gene expression**

Differential gene expression (DGE) analysis was performed using a custom version of the nextflow pipeline based on nf-core/differentialabundance (https://github.com/nf-core/differentialabundance) v.1.5.0 of the nf-core collection of workflows[1] using default parameters, including differential_max_qval = 0.05, with exceptions listed below. As input, salmon.merged.gene_counts.tsv and salmon.merged.gene_lengths.tsv were taken. Additionally, annotation assembly of the GRCh38 reference genome using Gencode v43[5] was used. Differentially expressed genes (DEGs) were defined using differential_min_fold_change = 1.5. DGE was performed by means of DESeq2[8] v.1.34.0. Data for each model and timepoint were analyzed separately using a simple contrast defining treatment with reference DMSO.

**Synergistically expressed genes**

Synergistically expressed genes (SEGs) were identified based on previous studies[9,10] with modifications. The analysis was performed using publicly available packages including *dplyr*[11] and others from the *tidyverse* collection, *edgeR* for CPM calculation[12] and *pheatmap* (https://github.com/raivokolde/pheatmap) for visualization. Lists of DEGs were obtained as described above. Only genes identified as DEGs in at least one comparison were considered for SEG analysis. In short, SEG analysis assigns predefined labels consisting of three parts: (i) UP/DN: the direction of expression change vs DMSO control; (ii) treatment label: indicates in which treatment (romaciclib, VEN, or RVU120+VEN) a gene was identified as a DEG; (iii) MORE label: indicates the presence of a synergistically driven gene expression change. This part of the label is based on the ratio of expression change (R) calculated according to a formula: $R = |COMBO - DMSO| / (|RVU120 - DMSO| + |VEN - DMSO|)$, where counts were obtained from nf-core rnaseq pipeline output files and transformed to CPM. $R > 1.25$ suggests a synergistic effect ("MORE" label). If a gene is a DEG only in the RVU120+VEN combination, it is automatically labelled as "MORE." These labels are assigned with an assumption that in all treatments the direction of change was identical. In case of mixed direction of expression change, such cases were labelled as "MIX."

**Weighted gene co-expression analysis**

Gene counts from processing of RNA-seq data were filtered, keeping genes having at least 15 counts in at least 75% of samples in the analyzed dataset. Variance stabilizing transformation was performed from DESeq2[8] v.1.38.3. Binary traits for each treatment were prepared. An optimal power for signed network was selected based on the results obtained using the pickSoftThreshold function from Weighted gene co-expression analysis (WGCNA)[13] v.1.71. Automatic network construction and module detection was done using blockwiseModules from WGCNA with default parameters except power (= 26, PDXAMLX12 and = 30, KG-1 3d), maxBlockSize = 23000, networkType = "signed," min_module = 30, reassignThreshold = 0, mergeCutHeight = 0.25 (= 0.2, KG-1 3d), numericLabels = TRUE, pamRespectsDendro = FALSE, randomSeed = 1234. Module-trait correlation was calculated using cor function and Student asymptotic p-value for correlation using corPvalueStudent function from WGCNA v.1.71. Student asymptotic p-values for correlation were denoted as follows: ***p-value < 0.001, **p-value < 0.01 and *p-value < 0.05. Heatmaps presenting module-trait correlation were prepared with ComplexHeatmap[14] v.2.14.0.

**Gene set enrichment and over-representation analysis**

Gene set enrichment analysis (GSEA) was performed using gsea software[15] version 4.3.2. Parameters were set as follows: nperm = 1000, metric = log2_Ratio_of_Classes, set_max = 500, set_min = 15, permutation = gene_set, scoring_scheme = weighted, sort = real, order = descending, norm = meandiv, and rnd_type = no_balance. Over-representation analysis (ORA) was performed using the compareCluster function from clusterProfiler[16] v.4.6.2 and default parameters except pvalueCutoff = 0.1 and maxGSSize = 5000 (ORA on DEGs). For both analyses, gene sets were obtained from msigdbr v.7.5.1, and bubble plots presenting enrichment results were prepared using the dotplot function from enrichplot v.1.18.4.

### Supplemental Tables

Supplemental Tables 1-6 are provided as separate .xls files.

| | |
|---|---|
| **Supplemental Table S1** | Media composition and origin of cell lines. |
| **Supplemental Table S2** | Primary AML sample characteristics. |
| **Supplemental Table S3** | Antibodies used for western blotting. |
| **Supplemental Table S4** | Enrichment of H/C2 collection gene sets in SEGs downregulated by RVU120+VEN in MV4-11 and MOLM-13 cells. |
| **Supplemental Table S5** | Enrichment of H/C2 collection gene sets in WGCNA modules in KG-1 cells treated with RVU120 +/- VEN. |
| **Supplemental Table S6** | Enrichment of H/C2 collection gene sets in WGCNA modules in PDX-AMLX12 cells treated with RVU120 +/- VEN. |

**Supp Fig.1**

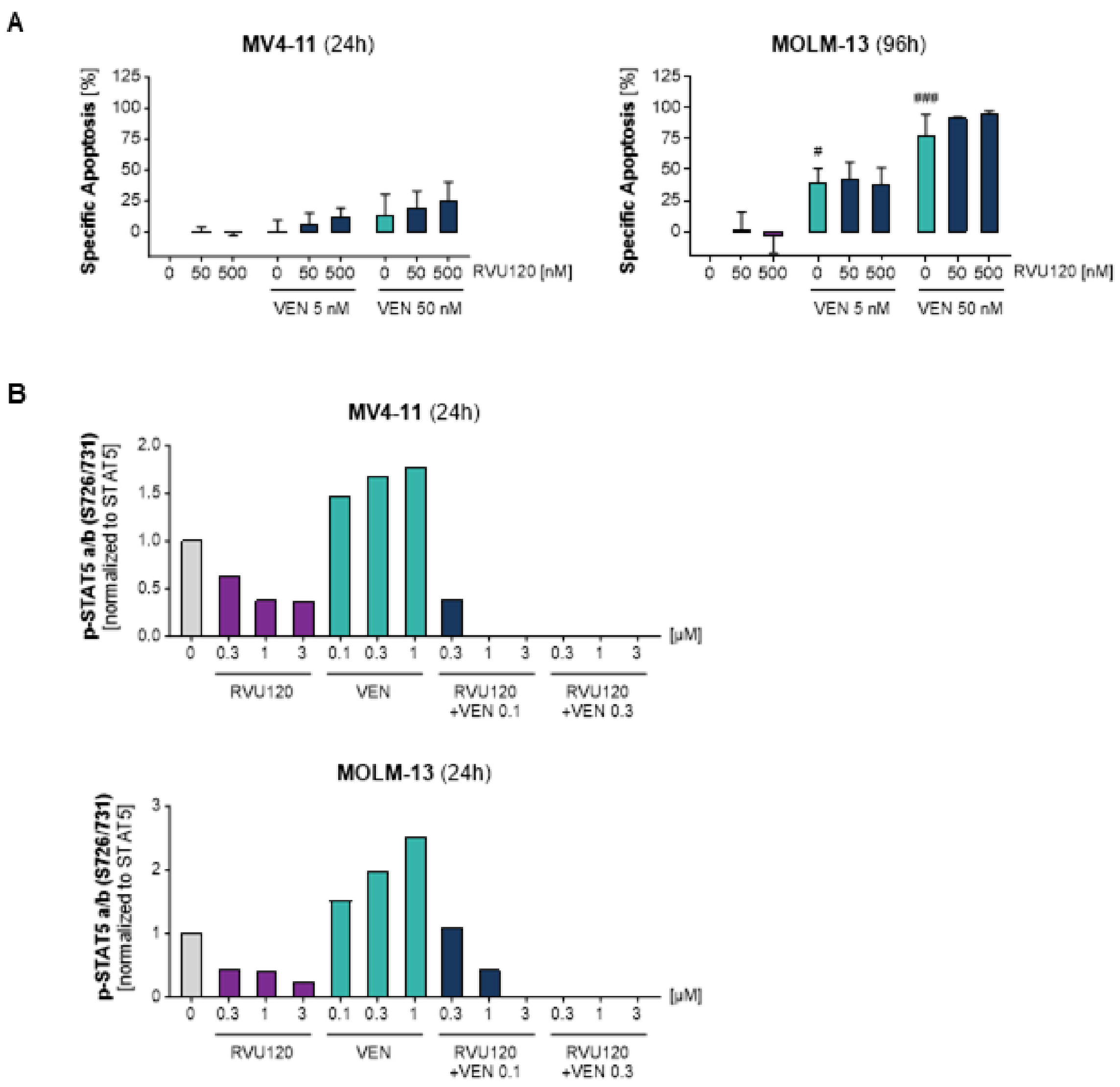


**Supplemental Figure S1. Romaciclib synergistically enhances antileukemic activity of VEN in VEN-sensitive AML cells.**

**(A**) Apoptosis induction in MV4-11 and MOLM-13 cells treated with romaciclib (50, 500 nM), VEN (5, 50 nM), or their combination at the indicated time points. Data are mean ± SD of two independent experiments. Two-way ANOVA with Tukey's multiple comparisons: $^{\#}$p<0.05, $^{\#\#\#}$p<0.001 (vs vehicle control). **(B)** Quantification of p-STAT5 (S726/731) level in in MV4-11 and MOLM-13 cells treated with the indicated concentrations of romaciclib and/or VEN for 24 hours. Data are representative for two independent experiments except for 0.1uM VEN concentration.

**Supp Fig.2**

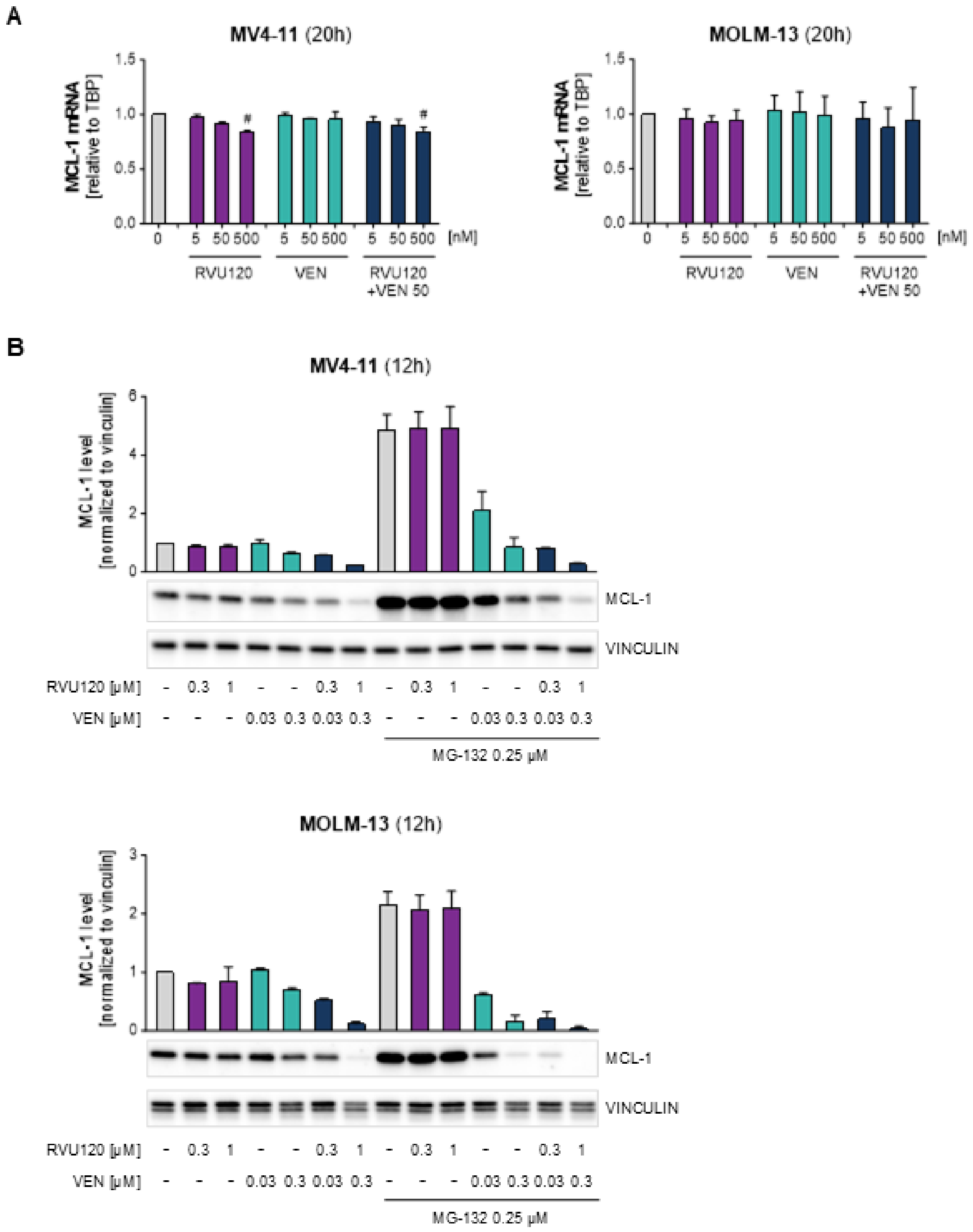


**Supplemental Figure S2. MCL-1 protein downregulation by RVU120+VEN is independent of transcriptional regulation and proteasomal degradation.**

**(**A) Relative mRNA expression of MCL-1 in MV4-11 and MOLM-13 cells after treatment with indicated concentrations of romaciclib and/or VEN for 20 hours. Data are presented as mean ± SD of two independent experiments comprising three technical replicates each. Two-way ANOVA with Tukey's multiple comparisons: $^{\#}p < 0.05$ (vs vehicle control). **(B)** Protein expression of MCL-1 in MV4-11 and MOLM-13 cells exposed to the indicated concentrations of romaciclib and/or VEN for 12 hours, with or without the addition of proteasome inhibitor MG-132 (12 hours, 0.25 µM). The bar graphs show quantification of MCL-1 protein levels. Representative western blots of two independent experiments are shown.

Supp Fig.3

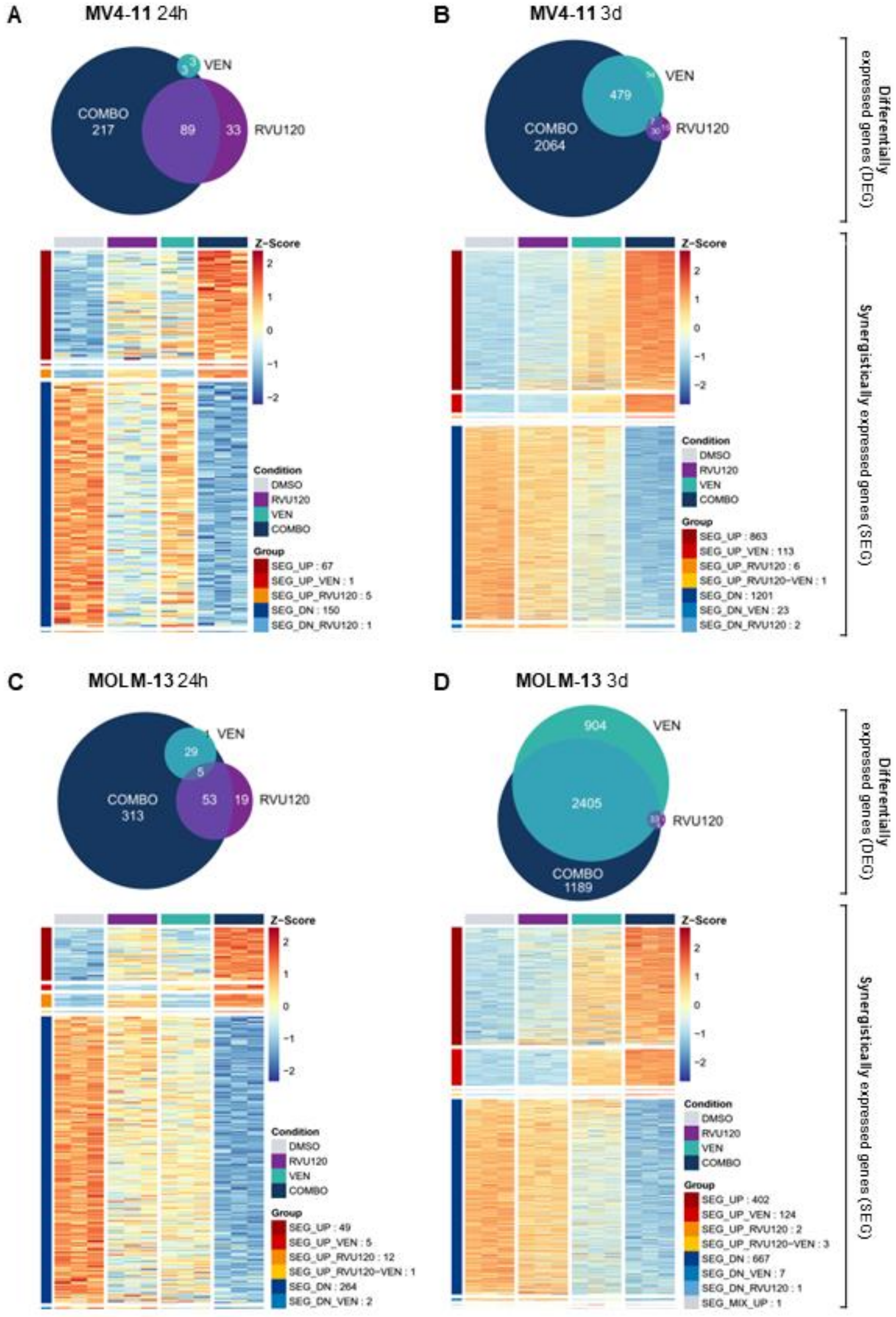
A
MV4-11 24h
VEN
COMBO
217
89
33
RVU120
Z-Score
Condition
DMSO
RVU120
VEN
COMBO
Group
SEG_UP : 67
SEG_UP_VEN : 1
SEG_UP_RVU120 : 5
SEG_DN : 150
SEG_DN_RVU120 : 1
B
MV4-11 3d
VEN
479
COMBO
2064
RVU120
Z-Score
Condition
DMSO
RVU120
VEN
COMBO
Group
SEG_UP : 863
SEG_UP_VEN : 113
SEG_UP_RVU120 : 6
SEG_UP_RVU120-VEN : 1
SEG_DN : 1201
SEG_DN_VEN : 23
SEG_DN_RVU120 : 2
Differentially expressed genes (DEG)
Synergistically expressed genes (SEG)
C
MOLM-13 24h
VEN
29
5
COMBO
313
53
19
RVU120
Z-Score
Condition
DMSO
RVU120
VEN
COMBO
Group
SEG_UP : 49
SEG_UP_VEN : 5
SEG_UP_RVU120 : 12
SEG_UP_RVU120-VEN : 1
SEG_DN : 264
SEG_DN_VEN : 2
D
MOLM-13 3d
904
VEN
2405
RVU120
COMBO
1189
Z-Score
Condition
DMSO
RVU120
VEN
COMBO
Group
SEG_UP : 402
SEG_UP_VEN : 124
SEG_UP_RVU120 : 2
SEG_UP_RVU120-VEN : 3
SEG_DN : 667
SEG_DN_VEN : 7
SEG_DN_RVU120 : 1
SEG_MIX_UP : 1
Differentially expressed genes (DEG)
Synergistically expressed genes (SEG)

**Supplemental Figure S3. RVU120+VEN combination drives synergistic gene expression changes in MV4-11 and MOLM-13 cells.**

Euler plots showing differentially expressed gene (DEG) counts and heatmaps showing Z-scores for synergistically expressed genes (SEGs) following treatment with romaciclib (100 nM), VEN (10 nM), or their combination in **(A)** MV4-11 at 24 hours, **(B)** MV4-11 at 3 days, **(C)** MOLM-13 at 24 hours, and **(D)** MOLM-13 at 3 days.

**Supp Fig.4**

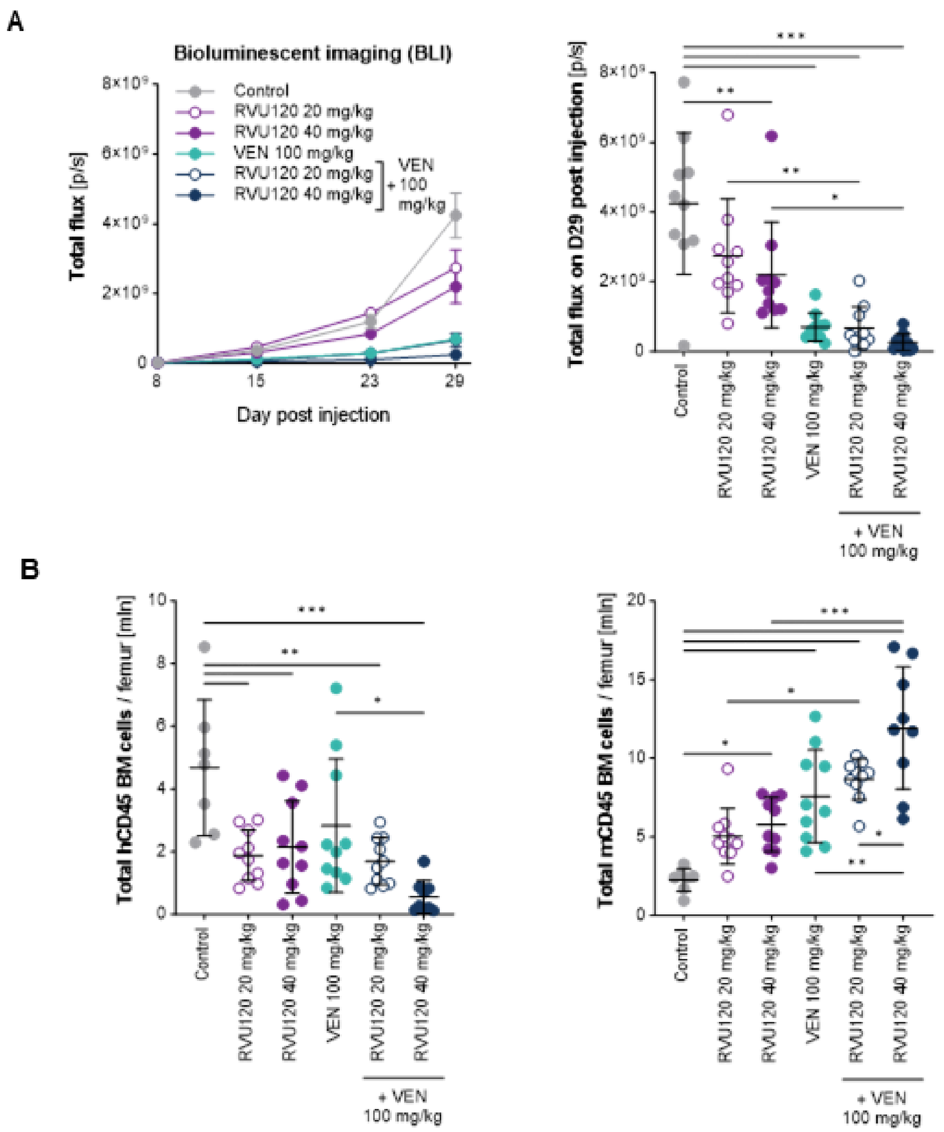


**Supplemental Figure S4. Preclinical in vivo efficacy of RVU120+VEN combination in orthotopic Luc-MV4-11 xenograft model.**

**(A)** Summary of BLI (Bioluminescence Imaging) changes over time for the treatment groups of mice in the study. Data are presented as mean ± SEM. BLI measurements of individual mice on day 29 post injection. Data are presented as mean ± SD. One-way ANOVA with Tukey's multiple comparisons: $^{*}p < 0.05$, $^{**}p < 0.01$, $^{***}p < 0.001$ (as indicated). **(B)** Quantification of human (hCD45) and murine (mCD45) bone marrow cells per femur after 21 days of treatment. Each data point represents an individual mouse. Data are presented as mean ± SD. One-way ANOVA with Tukey's multiple comparisons: $^{*}p < 0.05$, $^{**}p < 0.01$, $^{***}p < 0.001$ (as indicated).

**Supp Fig.5**

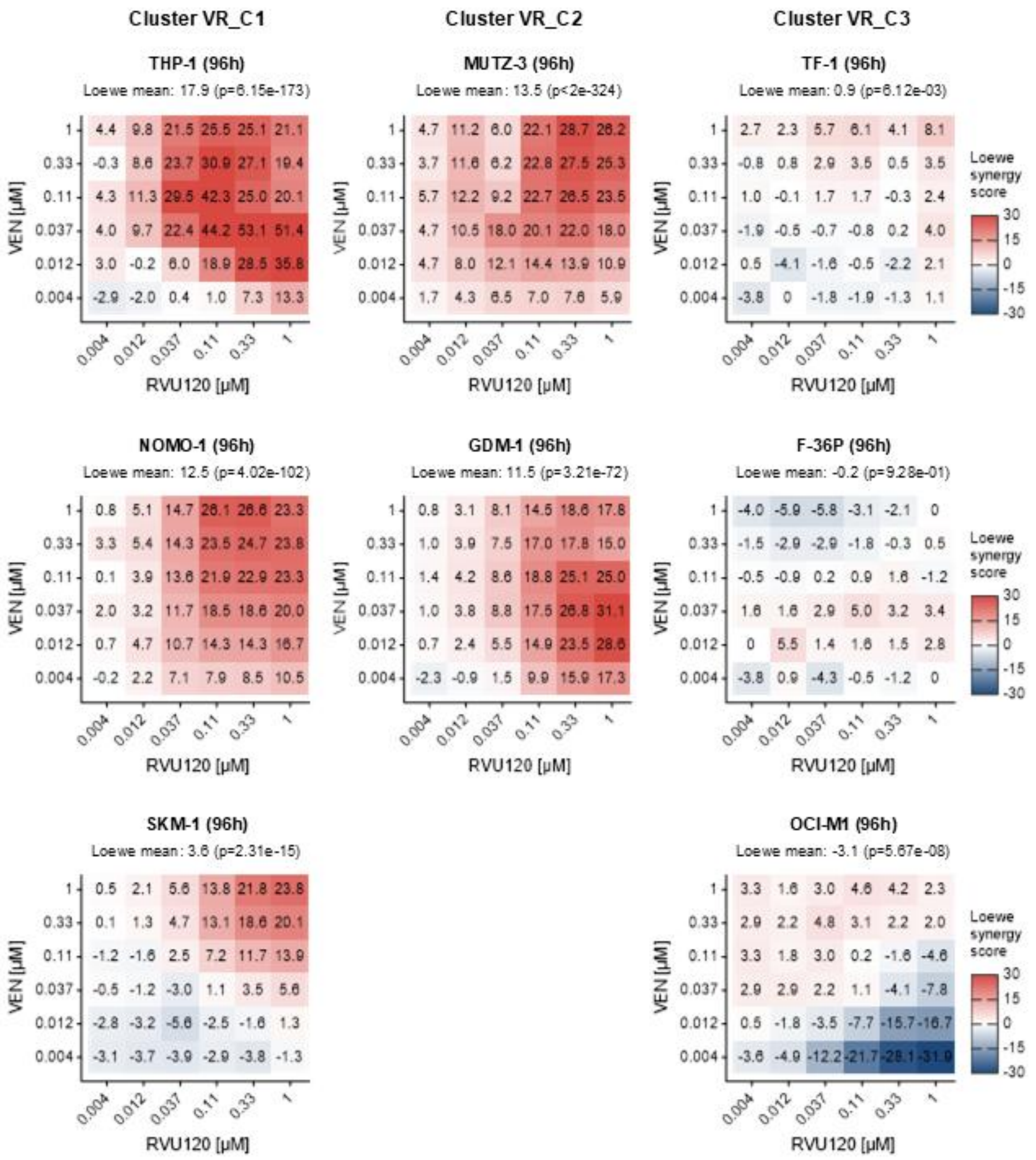


**Supplemental Figure S5. Romaciclib and VEN demonstrate synergistic antileukemic activity in VEN-resistant AML cells.**

Loewe synergy scores calculated from the effect of RVU120+VEN combination on viability of VEN resistant AML cell lines exposed to the indicated concentrations of drugs for 4 days. Loewe scores of > 0 are regarded as synergistic. Data of three technical replicates of representative independent experiment are presented (n = 2).

**Supp Fig.6**

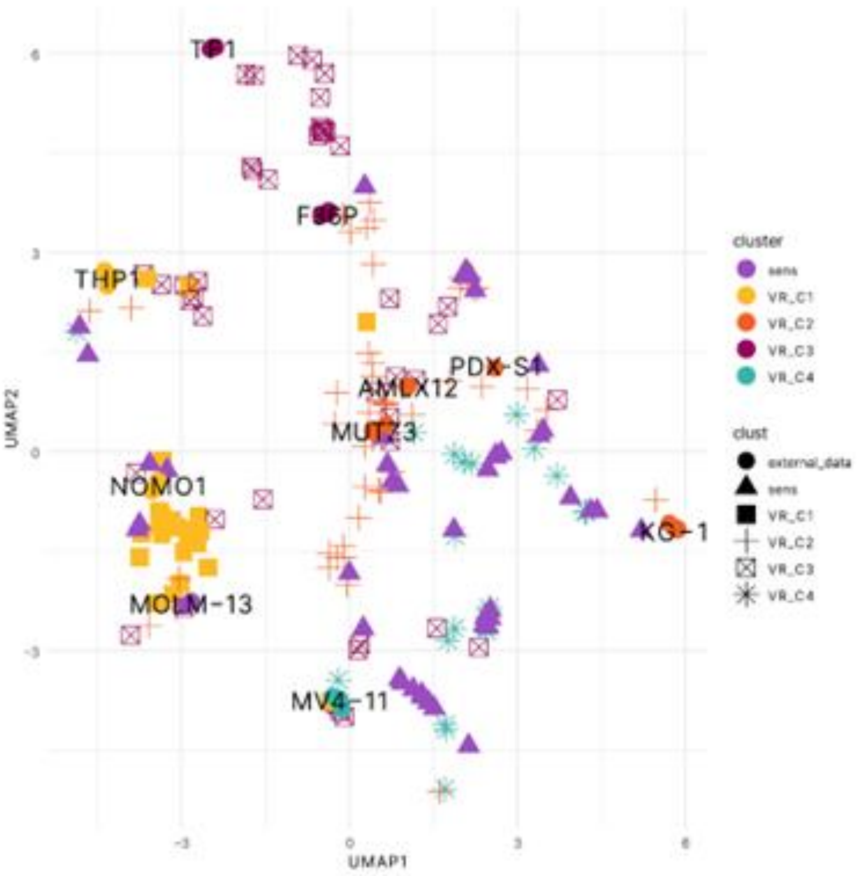


**Supplemental Figure S6. UMAP analysis of AML cell lines, PDX AMLX12 and PDX-S1.**

UMAP (Uniform Manifold Approximation and Projection) plot visualizes the clustering of AML cells based on their transcriptional profiles. Each point represents an individual cell line, PDX AMLX12 cells, or PDX-S1 cells with color coding indicating distinct clusters.
